# The nucleolar complex FAN-FIP1 mediates ribosome biogenesis in Arabidopsis and is critical for BR signaling and heat tolerance

**DOI:** 10.64898/2026.06.17.732803

**Authors:** Ya-Nan Wu, Jin-Yu Lu, Yang Gao, Sha Li, Feng Xiong, Yan Zhang

## Abstract

Ribosome biogenesis is critical for plant development and environmental responses. A large number of ribosomal proteins (RPs) and ribosomal biogenesis factors (RBFs) are required for ribosome biogenesis, many of which remain uncharacterized in plants. We report here the identification of Arabidopsis RBF FAN and its interacting partner FAN-INTERACTING PROTEIN 1 (FIP1). As their human and yeast orthologues, FAN-FIP1 interact. Both FAN and FIP1 participate in the processing of pre-rRNAs. Functional loss of *FAN* or *FIP1* knock-down results in developmental retardation and hypersensitivity to heat stresses. We demonstrate that FAN-FIP1 positively mediates brassinosteroid (BR) signaling by ensuring the translation efficiency of the BR receptor-coding gene *BRASSINOSTEROID INSENSITIVE 1* (*BRI1*) through the presence of its upstream open reading frame (uORF). Importantly, BR signaling positively mediates the processing of pre-rRNAs, which may be critical not only for development but also for heat tolerance.

## Introduction

Ribosome biogenesis is crucial for cell growth, differentiation, and division. In eukaryotes, ribosome biogenesis involves the processing of pre-ribosomal RNAs (pre-rRNAs) and sequential assembly of a large number of ribosomal proteins (RPs) on the rRNAs (Granneman & Baserga, 2004, Weis et al., 2015). In Arabidopsis, ribosome biogenesis begins with the transcription of the 45S rDNA gene to produce a 45S pre-rRNA primary transcript, which is eventually processed as 18S, 5.8S, and 25S rRNAs (Weis et al., 2015, Sáez-Vásquez & Delseny, 2019). The 18S rRNA and approximately 33 RPs are assembled into the 40S subunit whereas 5.8S, 25S, and 5S rRNAs assemble with approximately 47 RPs to form the 60S subunit (Weis et al., 2015, Sáez-Vásquez & Delseny, 2019). In the past two decades, functional studies on RP-coding genes in Arabidopsis have demonstrated that RPs mediated a full spectrum of developmental processes, such as shoot growth, root growth, leaf development (Horiguchi et al., 2011), embryogenesis (Szakonyi & Byrne, 2011), and gametophytic development (Byrne, 2009, Horiguchi et al., 2012).

In additional to RPs, ribosome biogenesis requires ribosome biogenesis factors (RBFs) (Weis et al., 2015), which transiently associate with pre-ribosomal particles and function as chaperones or as modification, processing, assembly, or remodeling factors (Dörner et al., 2023). Approximately 250 RBFs have been identified in the yeast genome (Henras et al., 2008), for which orthologues of 179 have been identified in plants (Ebersberger et al., 2014, Simm et al., 2015). Similar to mutations at RPs in plants (Byrne, 2009, Horiguchi et al., 2012), those of RBFs often result in pleiotropic phenotypes, such as impaired root growth (Wieckowski & Schiefelbein, 2012), abnormal cotyledons (Weis et al., 2014), defective gametophyte development (Shi et al., 2005, Li et al., 2009, Huang et al., 2010, Liu et al., 2010), and flowering delay (Lange et al., 2011, Weis et al., 2015). Despite these progresses, the biological and molecular function of most RBFs in plants are unclear.

AATF/Che-1 is a crucial RBF in metazoans (Kaiser et al., 2020). AATF/Che-1 interacts with 45S pre-rRNA, snoRNAs, rRNA processing proteins, RPs, and other RBFs to mediate the biogenesis of the 40S ribosomal subunit (Bammert et al., 2016, Kaiser et al., 2019). AATF/Che-1 also interacts with RNA polymerase I machinery to mediate rDNA transcription (Sorino et al., 2020). Mutations of a AATF/Che-1 homolog in mouse result in embryo arrest likely from compromised ribosome biogenesis (Thomas et al., 2000). The yeast homolog of AATF, Brefeldin A resistance protein 2 (BFR2), also participates in rRNA processing and RNA splicing as part of the yeast small subunit (SSU) processome (Soltanieh et al., 2014), suggesting that function of AATF/Che-1 is evolutionarily conserved. However, whether there is a similar mechanism in plants remains to be resolved.

The orthologue of yeast BFR2 and metazoan AATF/Che-1 in Arabidopsis is named FAN (Liu et al., 2023). FAN was recently reported to mediate the maintenance of root apical meristem (RAM) upon DNA damage through ATM/RAD53-RELATED (ATR) pathway (Liu et al., 2023). Whether FAN plays a role in ribosome biogenesis, as its orthologues in other phyla, is unclear. We report here that FAN and its interacting partner FIP1 mediate ribosome biogenesis by facilitating the processing of pre-rRNAs. We further show that FAN-FIP1 positively mediates brassinosteroid (BR) signaling by ensuring the translation efficiency of the BR receptor-coding gene *BRASSINOSTEROID INSENSITIVE 1* (*BRI1*) through the presence of its upstream open reading frame (uORF). Importantly, BR signaling positively mediates the processing of pre-rRNAs, which may be critical for plant development and stress responses.

## Results

### *fan* is defective in ribosome biogenesis and compromised in cell cycle progression

To test whether Arabidopsis FAN played a role in ribosome biogenesis, we first performed polysome profiling assays (Xiong et al., 2020). Compared with those in wild type, *fan* contained a substantial reduction in 40S and 60S subunits, as well as 80S and polysome fractions (Fig. 1a), similar to that in the mutants of RBF-coding genes (Cho et al., 2013, Choi et al., 2020), a defect fully rescued by *FANg:GFP* (Fig. 1a). To verify the reduction of ribosome biogenesis in *fan*, we introduced a two-expression-cassette *35S:ARF3-mRFP*;*35S:uORF-ARF3-GFP* (Fig. 1b) (Xiong et al., 2020) into the protoplasts of wild-type and *fan* mutants, respectively. Translational capacity of *AUXIN RESPONSE FACTOR3* (*ARF3*) was significantly compromised by an upstream open reading frames (uORFs) located in its 5’UTR preceding the main ORF (mORF), especially in defective mutants of ribosome biogenesis (Rosado et al., 2012). By comparing the fluorescence intensity of ARF3-GFP and ARF3-mRFP (Fig. 1c), we determined that the ratio of mRFP/GFP was significantly higher in *fan* than that in wild type (Fig. 1d), further supporting that ribosome biogenesis was compromised in *fan*.

**Figure 1.**
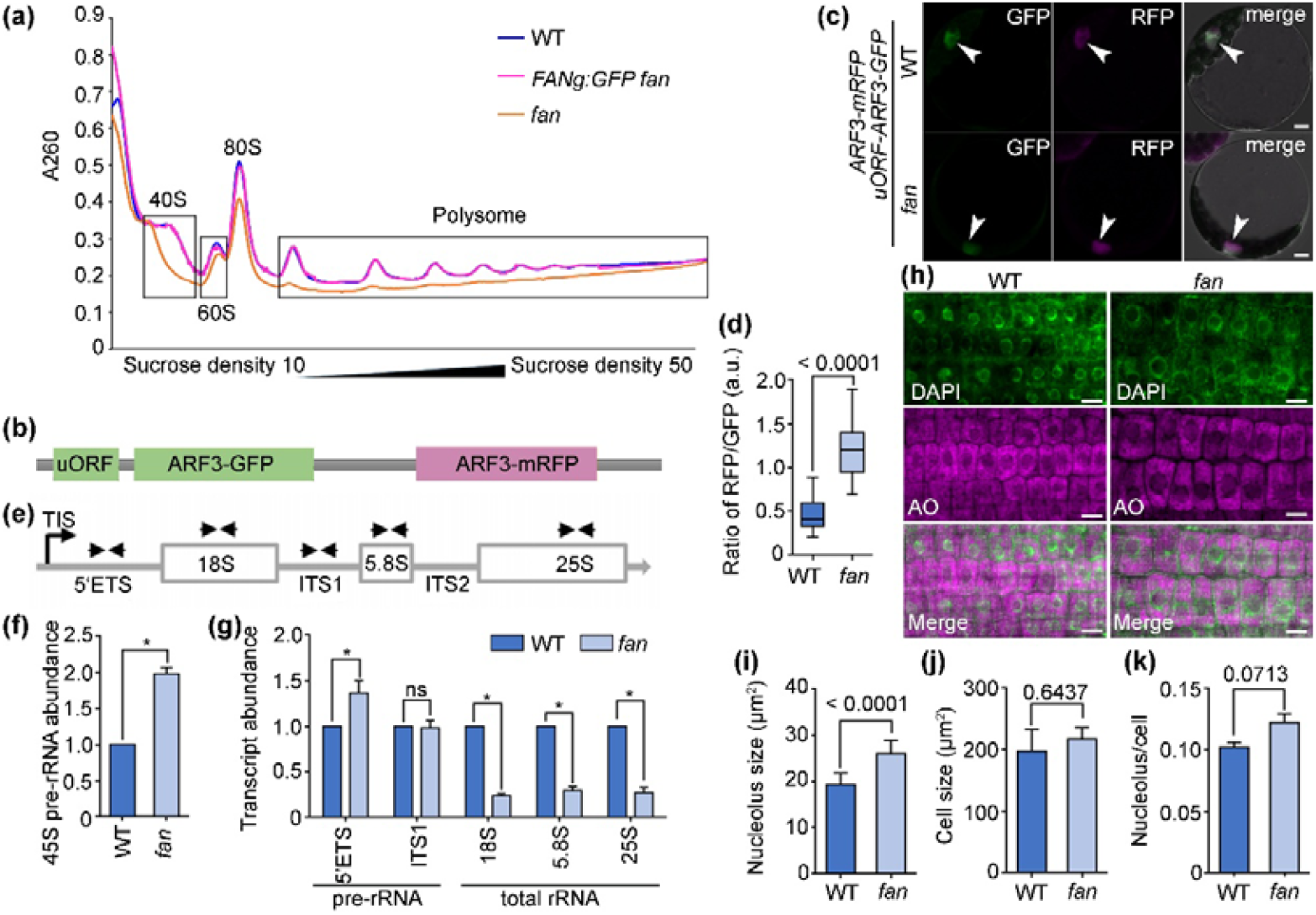
*fan* is defective in ribosome biogenesis. (a) Polysome profiling assay with sucrose density gradient. The OD_260_ absorption (A260) was monitored together with fractionation. The fractions containing 40S, 60S, 80S of ribosomes, and polysomes in wild type (WT), *FANg:GFP fan*, or *fan* are indicated. Three biological replicates were performed with similar results. (b) A cartoon illustrating the two-expression-cassette vector (*35S:ARF3-mRFP;35S:uORF-ARF3-GFP*) used in the protoplast expression assays. (c) Representative confocal laser scanning microscopic (CLSM) of a wild-type or *fan* protoplast transformed with *35S:ARF3-mRFP;35S:uORF-ARF3-GFP*. Merges of the GFP, RFP, and transmission channels are shown at the right. Arrowheads point at the nuclei. (d) Ratio of RFP versus GFP intensity in the nucleus from wild-type or *fan* protoplasts transformed with *35S:ARF3-mRFP;35S:uORF-ARF3-GFP*. Results are means ± standard deviation (SD, n > 30) from two batches of transformation events. For boxplots, the boxes extend from the 25^th^ to 75^th^ percentiles and the lines in the middle of the boxes are plotted at the median. P value is shown on top (*t*-test). (e) Representation of 45S pre-rRNA indicating positions of primer pairs used to amplify fragments containing the 5’ETS, ITS1, 18S, 5.8S, or 25S regions. ETS: external transcribed spacer; ITS: internal transcribed spacer. (f-g) RT-qPCRs of 45S pre-rRNA (f) or rRNA fragments (g) in wild type or *fan* seedlings at 7 d after germination (DAG). Expression levels are related to that of *EF-1α* (left) or 45S pre-rRNA (right). Results are means ± standard errors (SE, n = 3). Asterisks indicate significant difference (*t*-test, P < 0.05). (h) CLSM of DAPI and acridine orange (AO) staining from wild-type or *fan* roots at 5 DAG. DAPI (green) was used to label cell nucleus, AO (magenta) was used to label nucleolus. (i-k) Nucleolus size (i), cell size (j) or ratio of nucleolus size versus cell size (k) from wild-type or *fan* roots at 5 DAG. Results are means ± SD (n > 30). P values are shown on top (*t*-test). Bars = 5 µm (d); 10 µm (h).

To determine at which steps during ribosome biogenesis *fan* was defective, we examined the levels of pre-rRNAs and mature 5.8S, 18S, and 25S rRNAs by reverse-transcription quantitative PCRs (RT-qPCRs) (Fig. 1e). To exclude the influences of transcriptional differences between wild type and *fan*, we examined the expression of 45S precursor (Fig. 1f) and used the levels of 45S pre-rRNAs for normalization as reported (Chen et al., 2016). Consistent with a defective ribosomal biogenesis as shown by ribosomal profiling, the levels of pre-rRNAs in *fan* were significantly higher than those in wild type (Fig. 1g). By contrast, the levels of total rRNAs in *fan* were significantly lower than those in wild type (Fig. 1g). Introducing *FANg:GFP* in *fan* fully rescued its defective pre-rRNA processing as indicated by the levels of 45S rRNA, pre-rRNAs and total rRNAs (Fig. S1). These results suggest that *fan* is defective in the processing of pre-rRNAs, which would have caused a reduction in ribosomal biogenesis.

A reduced ribosome biogenesis often results in nucleolar hypertrophy (Micol-Ponce et al., 2018). We thus examined whether it was the case in *fan* by using a double labeling assay with both 4’,6-diamidino-2-phenylindole (DAPI) and acridine orange (AO), which emits fluorescence when bound to DNA or RNA, respectively (Micol-Ponce et al., 2018). Indeed, the size of nucleolus was significantly increased in *fan* compared to that in wild type (Fig. 1h-k), further supporting a reduced ribosome biogenesis in *fan*.

Reduced ribosome biogenesis often compromises cell cycle progression in yeast and metazoans (Dez & Tollervey, 2004, Donati et al., 2012). To determine whether it was the case in Arabidopsis, we examined cell ploidy by using flow cytometry. Indeed, *fan* contained more G1 (2C) cells and fewer G2 (4C) cells than wild-type (Fig. S2). In addition, a higher percentage of 16C and 32C cells was detected in *fan* than in wild type (Fig. S2), indicating that cells in *fan* tended to enter endo-replication. To further demonstrate the reduced cell cycle progression in *fan*, we introduced two cell cycle progression markers, *CycB1;1:GUS* and *KNOLLE:GFP* (González-García et al., 2011), into *fan* by crosses. By histochemical GUS staining or by confocal laser scanning microscopy (CLSM), we demonstrated that *fan* contained a substantial reduction of G2/M phase cells (Fig. S2), supporting a compromised cell cycle progression in *fan*.

### FAN interacts with FIP1 in the nucleolus

To identify interactors of FAN that contributed to pre-rRNA processing and ribosome biogenesis, we performed a genome-wide yeast two hybrid (Y2H) screening. Among the candidate interactors, we identified At1g07840, hereafter named FAN-INTERACTING PROTEIN 1 (FIP1). Arabidopsis FIP1 is homologous to human neuroguidin (NGDN) and yeast Lcp5 (Fig. S3), containing a Sas10 domain (Fig. 2a). Human NGDN interacts with AATF to form a nucleolar complex for 40S ribosomal subunit synthesis (Bammert et al., 2016) whereas yeast Lcp5 interacts with the yeast AATF orthologue Bfr2 to mediate pre-rRNA processing (Wiederkehr et al., 1998). All plant species contain FIP1 orthologues with the Sas10 domain (Fig. S3), suggesting a similar module composed of FAN-FIP1 for ribosome biosynthesis in plants.

**Figure 2.**
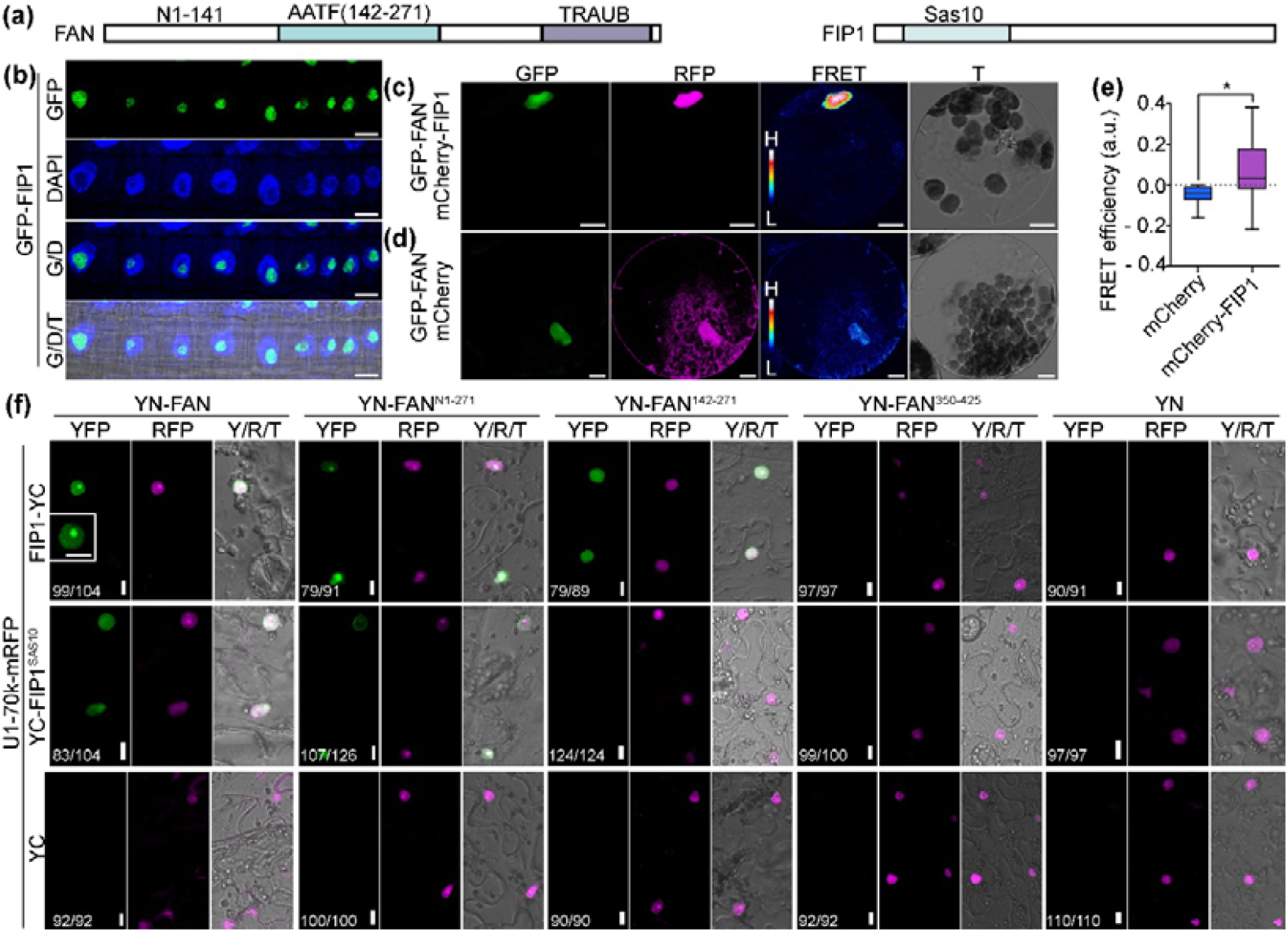
FAN interacts with a nucleolus protein FIP1. (a) Schematic illustration of FAN or FIP1 proteins used in this study. (b) CLSM images of root epidermal cells from the *UBQ10:GFP-FIP1* transgenic plants. DAPI staining was performed to indicate the nuclei. G/D indicates the merge of the GFP and DAPI channels; G/D/T indicates the merge of the GFP, DAPI, and transmission channels. (c-d) CLSM images of Arabidopsis leaf protoplasts expressing GFP-FAN/mCherry-FIP1 (c) or GFP-FAN/mCherry (d). FRET signals are represented in pseudocolors, covering the full range of measured values within each dataset (high to low). (e) Quantification of FRET efficiency. Results shown are means ± SD (n ≥ 15). Asterisk indicates significant difference (*t*-test, P < 0.05). (f) CLSM images of tobacco leaf epidermal cells in bimolecular fluorescence complementation (BiFC) assays. YN-FAN, YN-FAN^N1-271^, YN-FAN^142-271^ or YN-FAN^350-425^ was co-expressed with YC-fusions of FIP1 or FIP1^Sas10^, together with U1-70k-mRFP, a nucleus marker. Merge Y/R/T indicates merges of the YFP, RFP, and transmission channels. Numbers at the bottom indicate displayed (YFP-positive or negative cells) versus RFP-positive cells. Results are representative of three biological replicates. Bars = 10 µm.

Consistent with its potential role in ribosomal biogenesis, GFP-FIP1 was detected mostly in the nucleoli (Fig. 2b). *FIP1* was constitutively expressed in multiple tissues and developmental stages based on RT-qPCRs (Fig. S4). Consistent with RT-qPCRs, GUS signals from the *proFIP1:GUS* reporter lines were detected in most tissues and developmental stages tissues, such as seedlings, roots, rosette leaves, inflorescences, and ovules, as well as developing embryos (Fig. S4).

To verify the interaction between FAN and FIP1, we performed a Fluorescence Resonance Energy Transfer (FRET) assay, in which FIP1 and FAN colocalized and interacted (Fig. 2c-e). Next, we examined the interacting domains between FAN and FIP1 by bimolecular fluorescence complementation (BiFC). BiFC assays showed that FAN, FAN^N1-271^, or FAN^142-271^ interacted with FIP1 while FAN^350-425^ did not interact with FIP1 (Fig. 2f). On the other hand, FIP1^Sas10^ interacted with FAN or FAN^N1-271^, but not with FAN^142-271^ or FAN^350-425^ (Fig. 2f). To further verify the interacting domains between FAN and FIP1, we used Y2H assays. Similar to the results from BiFC, Y2H assays showed that the Sas10 domain within FIP1 and the AATF domain within FAN were key to their molecular interaction (Fig. S5). Molecular docking assays with AlphaFold3.0 and PyMOL further supported the involvement of FIP1-Sas10 domain and FAN-AATF domain in their interaction (Fig. S5).

To determine the biological relevance of FAN-FIP1 interaction, we introduced a *UBQ10*-driven GFP-translational fusion of *FIP1* (*UBQ10:GFP-FIP1*) construct into *fan*. By Western blot assay (Fig. 3a) and CLSM imaging (Fig. 3c-e), we determined that GFP-FIP1 was significantly reduced in *fan* as compared to that in wild type (Fig. 3a, 3c) despite that the transcript abundance of *GFP-FIP1* was comparable between two genotypes (Fig. 3b). Although GFP-FIP1 was distributed in the nucleus both in wild type (Fig. 3d) and in *fan* (Fig. 3e), a substantial reduction of GFP-FIP1 in the nucleolus of *fan* was detected (Fig. 3d-e).

**Figure 3.**
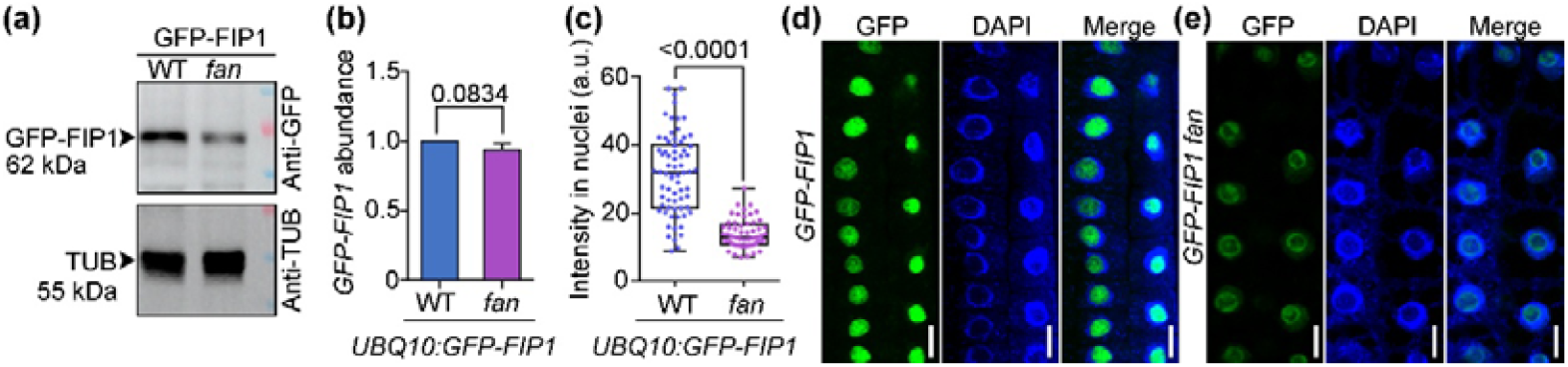
FAN facilitates the stabilization of FIP1. (a) A representative Western blot assay showing that GFP-FIP1 abundance in *UBQ10:GFP-FIP1* (WT) is substantially reduced in *UBQ10:GFP-FIP1 fan* (*fan*) seedlings at 7 DAG. With that in wild type being set as 1, the intensity of GFP-FIP1 in *fan* is 0.47 ± 0.05 (SD, n = 3). Ab-Tub (for tubulin) was used as the internal loading control. (b-c) Transcript abundance of *GFP-FIP1* (b) or intensity of nuclear GFP-FIP1 (c) in *UBQ10:GFP-FIP1* (WT) versus *UBQ10:GFP-FIP1 fan* (*fan*) seedlings at 5 DAG. (d-e) CLSM images of root epidermal cells from the *UBQ10:GFP-FIP1* or *UBQ10:GFP-FIP1 fan* seedlings at 5 DAG. Seedlings were stained with DAPI. Merges of the GFP and DAPI channel images are shown at the right side. Bars = 10 µm.

### FIP1 is critical for pre-rRNA processing and ribosome biogenesis

To test whether FIP1 participated in ribosomal biogenesis, we took a reverse genetic approach by examining T-DNA insertion lines at *FIP1*. None of the analyzed T-DNA insertion lines for *FIP1*, including SALK_136495, SALK_108967c, and SAIL_910_D01, germinated. We then tried to generate *FIP1* loss-of-function mutants by CRISPR/Cas9, which was also fruitless for unknown reasons. We then generated *FIP1* knock-down lines by using artificial microRNAs, which was demonstrated to be effective and genetically stable (Ossowski et al., 2008). We generated multiple transgenic plants with the expression vector *UBQ10:amiR-FIP1* and verified down-regulation of *FIP1* in two independent single-copy insertion lines by RT-qPCRs (Fig. 4a), which were used for phenotype analysis.

**Figure 4.**
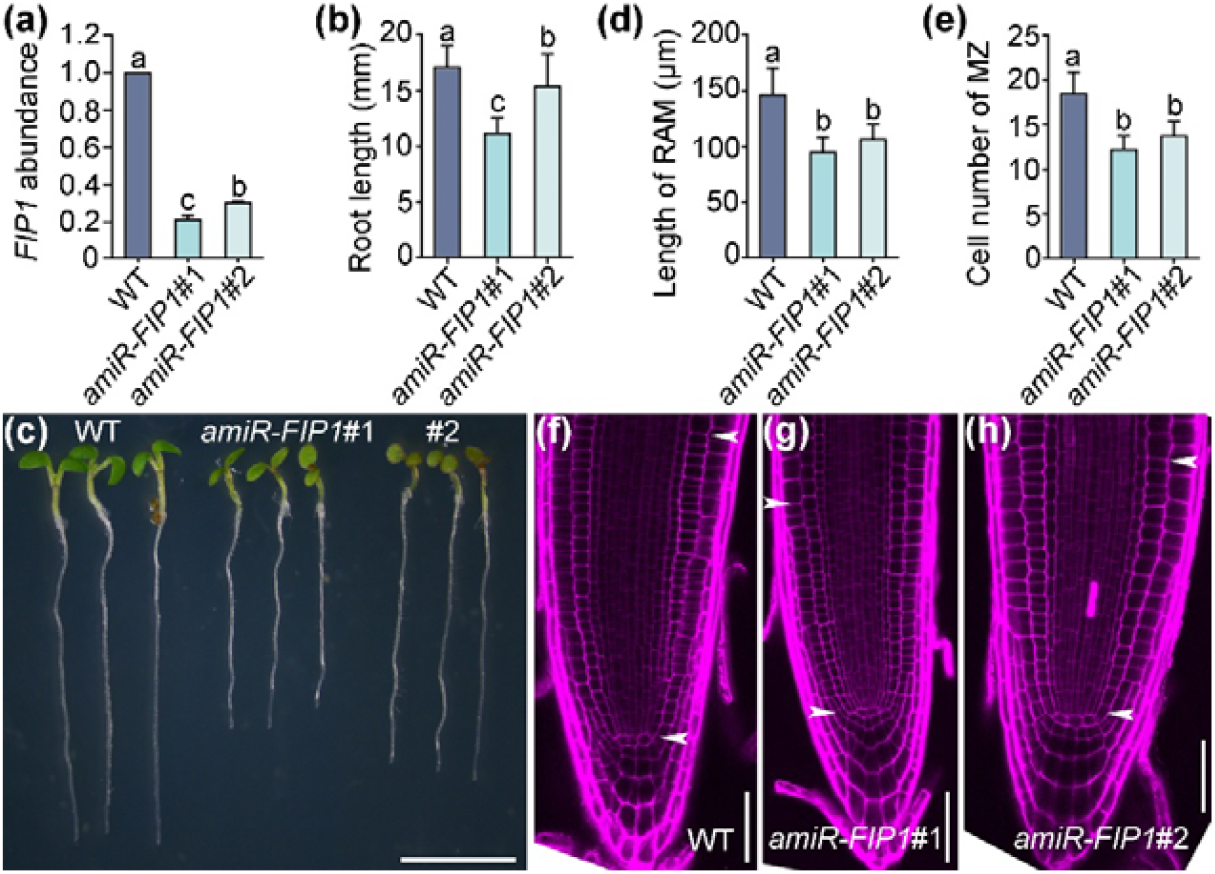
FIP1 knockdown compromises root growth. (a) Reverse transcription quantitative PCRs (RT-qPCRs) of *FIP1*. RNAs were extracted from roots of 7 DAG seedlings. Transcript abundance of *FIP1* in wild type was set as 1. Results shown are means ± SD (n = 3). Each biological replicates were repeated three times with similar results. (b) Primary root length from 7 DAG seedlings. Results are means ± SD (n > 20). (c) Representative seedlings at 7 DAG from wild type and two independent lines of *amiR-FIP1* transgenic plants. (d-e) Root apical meristem (RAM) length (d) or the number of meristematic cortex cells (MCC) in RAM (e) from 7 DAG seedlings. Results are means ± SD (n > 20). Different letters in (a, b, d, e) indicate significantly different groups (OneWay ANOVA, Tukey’s multiple comparisons test, P < 0.01). (f-h) CLSM of a PI-stained root tip from 7 DAG wild-type (f) and two independent lines of *amiR-FIP1* transgenic plants (g-h). RAM is between two arrows. Bars = 5 mm (c); 50 µm (f-h).

Knocking-down *FIP1* resulted in retarded growth (Fig. S6), similar to that of RP mutations (Horiguchi et al., 2012) and to that of *fan* (Liu et al., 2023). A close examination of root growth of *amiR-FIP1* lines showed that knocking-down *FIP1* significantly reduced root length (Fig. 4b-c). Because the development of root apical meristem (RAM) is crucial for root growth, we used PI staining to examine RAM development. Indeed, knocking-down *FIP1* significantly interfered with RAM development such that *amiR-FIP1* plants showed a significant reduction of RAM length (Fig. 4d, 4g-h) and of meristematic cortex cell (MCCs) number (Fig. 4e) compared to those of wild type (Fig. 4d-f), similar to that in *fan* (Liu et al., 2023).

To test whether FIP1 mediated ribosomal biogenesis, as its interacting partner FAN, we first examined *amiR-FIP1* transgenic plants in polysome profiling assays. Compared to those in wild type, knocking-down *FIP1* resulted in a substantial reduction in 40S and 60S subunits, as well as 80S and polysome fractions (Fig. 5a). To determine how knocking-down *FIP1* affected ribosome biogenesis, we examined the levels of pre-rRNAs and mature 5.8S, 18S, and 25S rRNAs by RT-qPCRs, after normalization with the levels of 45S precursor as for *fan* (Fig. 1f). Indeed, knocking-down *FIP1* resulted in significantly higher levels of pre-rRNAs but lower levels of total rRNAs (Fig. 5b-5c), indicating that knocking-down *FIP1* compromised the processing of rRNAs. Similar to *fan* (Fig. 1), *amiR-FIP1* plants contained relatively larger nucleoli than those in wild type (Fig. S7), further supporting its role in ribosomal biogenesis.

**Figure 5.**
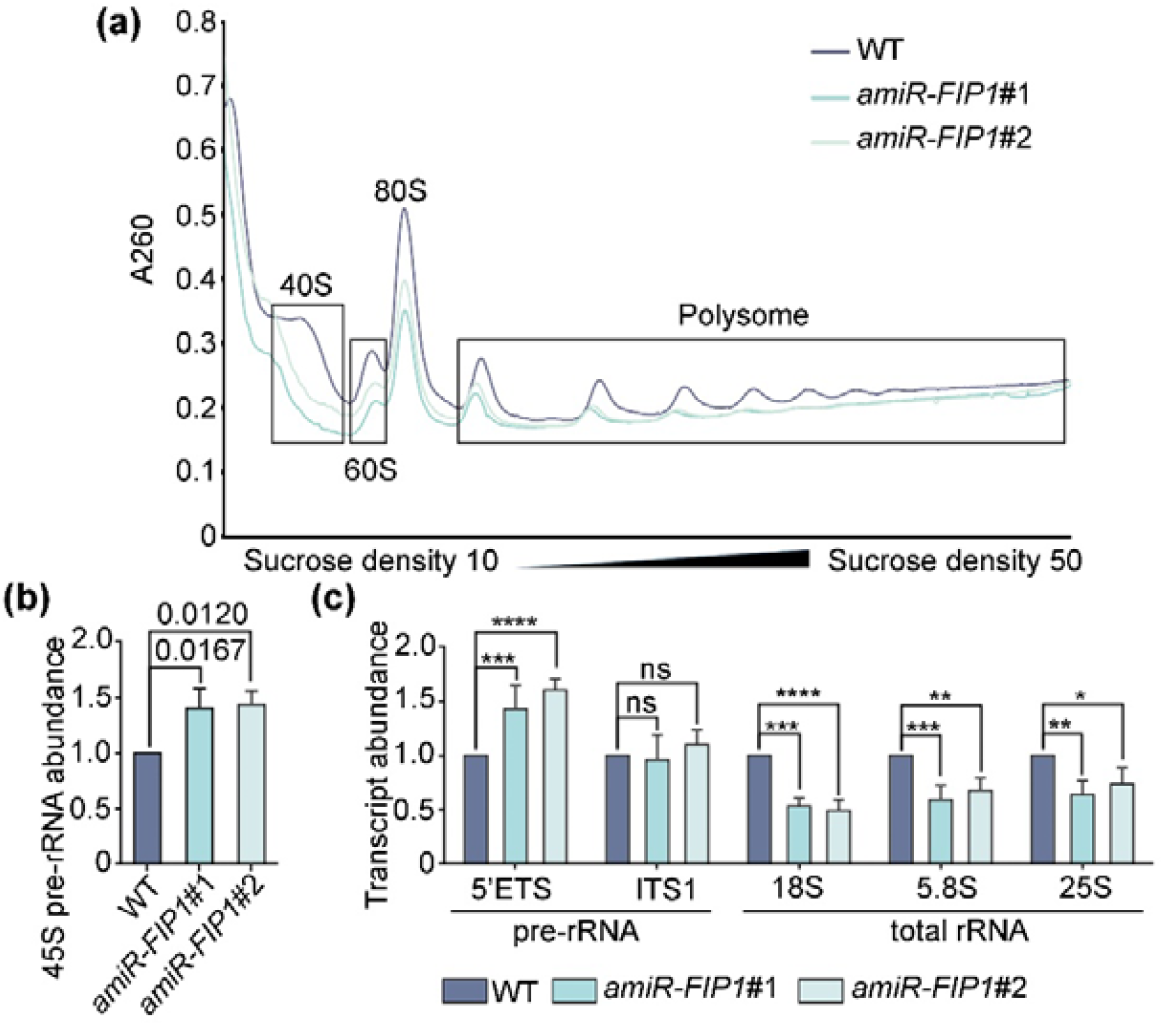
*FIP1* knockdown compromises ribosome biogenesis. (a) A representative polysome profiling assay with sucrose density gradient (the values 10 to 50 indicate the concentration of sucrose gradient). The OD_260_ absorption (A260) was monitored together with fractionation. The fractions containing 40S, 60S, 80S of ribosomes, and polysomes in wild type or two independent lines of *amiR-FIP1* are indicated. Three biological replicas were performed, showing similar patterns (reduction of ribosomal biogenesis in *amiR-FIP1* lines). (b-c) RT-qPCRs of 45S pre-rRNA (b) or rRNA fragments (c) abundance in wild type or two independent lines of *amiR-FIP1* roots from 7 DAG seedlings. Expression levels are related to that of *EF-1α* (left) or 45S pre-rRNA (right). Results shown are means ± SD (n = 3). Three biological replicates were performed with similar results. P values (*t*-test) for (b) are shown. For (c), * indicates P < 0.05; ** indicates P< 0.01; *** indicates P < 0.001; **** indicates P < 0.0001; ns indicates P > 0.05 (*t*-test).

### FAN-FIP1 positively mediates brassinosteroid signaling

Because FAN-FIP1 was critical for ribosomal biogenesis (this study) whereas brassinosteroid (BR) signaling pathway is sensitive to translation efficiency (Zhang et al., 2020), we wondered whether FAN-FIP1 mediated BR signaling.

To test this idea, we took a pharmacological approach by applying brassinolide (BL), the active analog of BR (Tang et al., 2008). Both *fan* and *amiR-FIP1* transgenic plants showed a reduced root growth without BL (Fig. 6a, 6e), as reported for *fan* (Liu et al., 2023). BL at 100 nM caused reduced root growth in wild type (Fig. 6a-b, 6e), as reported (Tang et al., 2008, Liu et al., 2024b). In comparison, root elongation of *fan* and *amiR-FIP1* transgenic plants showed a reduced sensitivity to BL (Fig. 6b, 6e). A reduced hypocotyl elongation in the dark in *fan* as well as *amiR-FIP1* transgenic plants (Fig. 6c, 6f) indicated reduced BR signaling. BL at 100 nM caused a reduction of hypocotyl elongation in wild type (Fig. 6c-d, 6f), as reported (Tang et al., 2008, Zhou et al., 2013b). In comparison, hypocotyl elongation of *fan* and *amiR-FIP1* transgenic plants showed a reduced sensitivity to BL (Fig. 6c-d, 6f).

**Figure 6.**
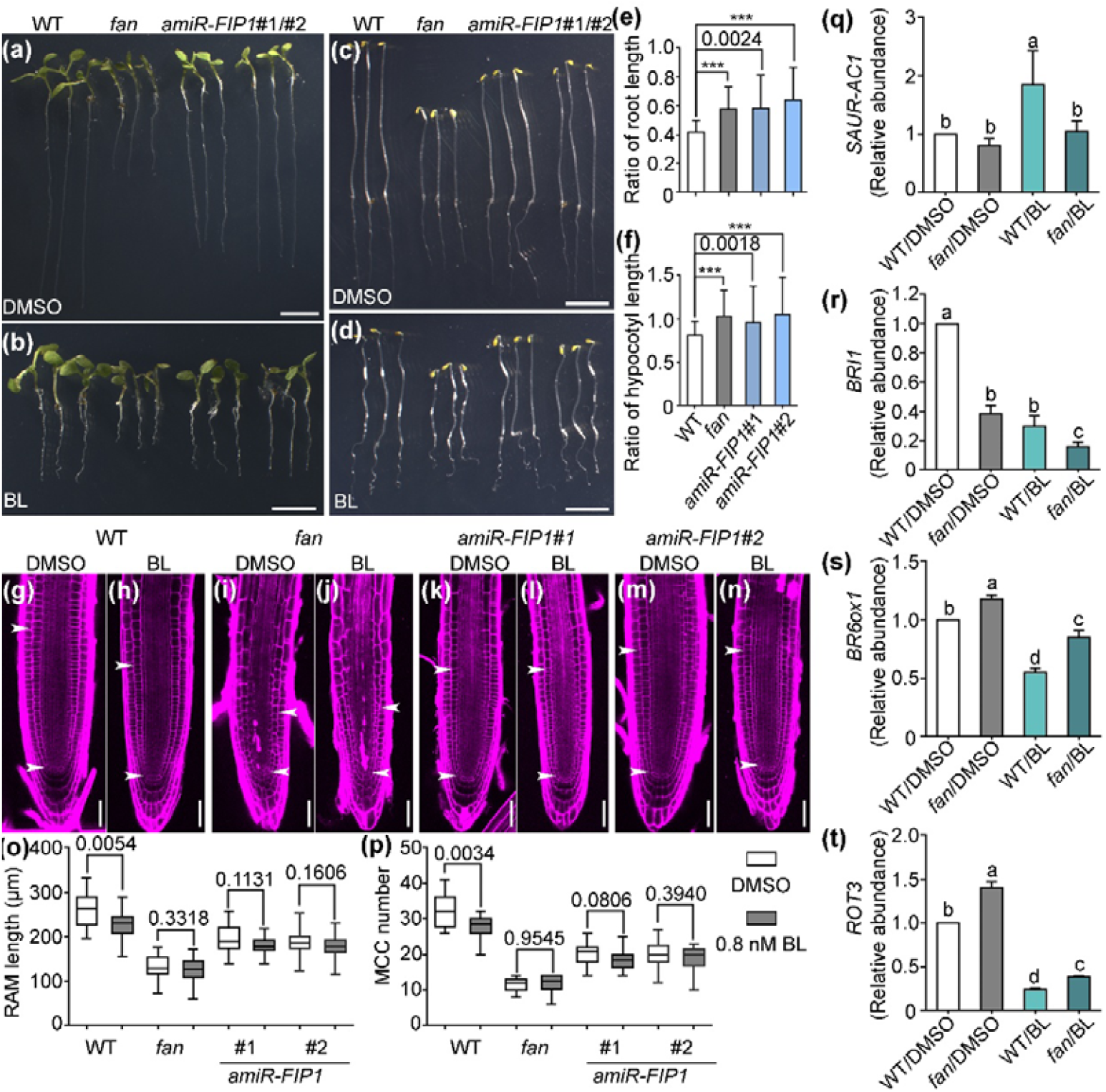
BL-responsive root development and hypocotyl elongation are compromised in *fan* or *amiR-FIP1* plants. (a-d) A root growth (a-b) or hypocotyl elongation assay (c-d) upon 100 nM BL treatment (b, d) versus DMSO (a, c). For root assay, sterilized seeds were grown on 1/2 MS plates supplemented with DMSO or 100 nM BL for 7 days; for hypocotyl assay, sterilized seeds were grown on 1/2 MS plates supplemented with DMSO or 100 nM BL for 5 days under dark. (e-f) 100 nM BL/DMSO ratio of root length (e) or hypocotyl length (f). Results shown are means ± SD (n > 15). (g-n) CLSM of a PI-stained root tip from 5 DAG seedlings of wild type (g-h), *fan* (i-j), and two lines of *amiR-FIP1* (k-n) treated with DMSO (g, i, k, m) or 0.8 nM BL (h, j, l, n). The region of RAM is indicated by two arrows. (o-p) RAM length (o) or the number of MCC in RAM (p) in seedlings supplemented with DMSO or 0.8 nM BL for 5 days. Results are means ± SD (n > 30). Asterisks (e-f) indicate P < 0.0001 or P values (e-f, o-p) are shown on top of the column (*t*-test). (q-t) RT-qPCRs of *SAUR-AC1* (q), *BRI1* (r), *BR6ox1* (s), or *ROT3* (t). RNAs were extracted from wild-type or *fan* seedlings as indicated in (a). Transcript abundance in DMSO-treated wild type was set as 1. Results shown are means ± SE (n = 3). Different letters indicate significantly different groups (OneWay ANOVA, Tukey’s multiple comparisons test, P < 0.01). Bars = 5 mm (a-d); 50 µm (g-n).

To provide further evidence that functional loss of *FAN* or knocking down *FIP1* compromised BR signaling, we examined the effect of BL on RAM. As reported (González-García et al., 2011), BL at 0.8 nM inhibited cell division at RAM in wild type as indicated by a reduced RAM length and a reduced number of meristematic cortex cells (MCC) (Fig. 6g-h, 6o, 6p). In comparison, both *fan* (Fig. 6i-j) and two independent *amiR-FIP1* transgenic plants (Fig. 6k-n) showed a reduced sensitivity to BL in regard to RAM development (Fig. 6o-p). These results support a reduced BR signaling by functional loss of *FAN* or knocking down *FIP1*.

Next, we examined the effect of exogenous BL application on the expression of BR signaling and biosynthetic genes by quantitative RT-PCRs. BL induced the expression of *SMALL AUXIN UP RNA 1* (*SAUR-AC1*) whereas repressed the expression of the BR receptor-coding gene *BRASSINOSTEROID INSENSITIVE 1* (*BRI1*) in wild type (Fig. 6q-r), as reported (Goda et al., 2002, Nakamura et al., 2003, Liu et al., 2024b). In comparison, *fan* seedlings were less responsive than wild type to the inductive or repressive effect of BL on *SAUR-AC1* (Fig. 6q) or *BRI1* (Fig. 6r), respectively. On the other hand, BL repressed the expression of BR biosynthetic genes *BRASSINOSTEROID-6-OXIDASE 1*/*CYTOCHROME P450 90B1* (*BR6ox1*) and *ROTUNDIFOLIA3*/*CYTOCHROME P450 90C1* (*ROT3*) in wild type (Fig. 6s-t), as reported (Tanaka et al., 2005). In comparison, the abundance of *BR6ox1* or *ROT3* was significantly higher in *fan* than in wild type without BL treatment (Fig. 6s-t). Exogenous application of BL caused a reduction of *BR6ox1* or *ROT3* in wild type (Fig. 6s-t). In comparison, *BR6ox1* in *fan* was less responsive to the inhibitory effect of BL (Fig. 6s), further supporting a compromised BR signaling by functional loss of *FAN*.

### FAN is important for maintaining translational efficiency of *BRI1*

Negative feedback on *BR6ox1* and *ROT3* transcription by BL relies on BRI1 (Tanaka et al., 2005) and *BRI1* contains a upstream open reading frames (uORF) in its 5′-UTR (Zhang et al., 2018, Zhang et al., 2020) that renders it sensitive to a decreased ribosome abundance (Zhang et al., 2019). We thus hypothesized that the reduced BR signaling in *fan* and *amiR-FIP1* was due to a reduced translation efficiency of *BRI1*.

To test this idea, we generated *UBQ10*-driven GFP-translational fusion *BRI1* expression constructs, with or without its endogenous uORF, i.e. *UBQ10:uORF-BRI1-GFP* or *UBQ10:BRI1-GFP*. The constitutive *UBQ10* promoter was used to exclude the influence of *BRI1* transcription. We obtained single-copy insertional *UBQ10:uORF-BRI1-GFP* and *UBQ10:BRI1-GFP* transgenic plants in wild type and crossed the transgenes into *fan* so that the same transgene was used for comparison between wild type and *fan* (Fig. 7a, 7e).

**Figure 7.**
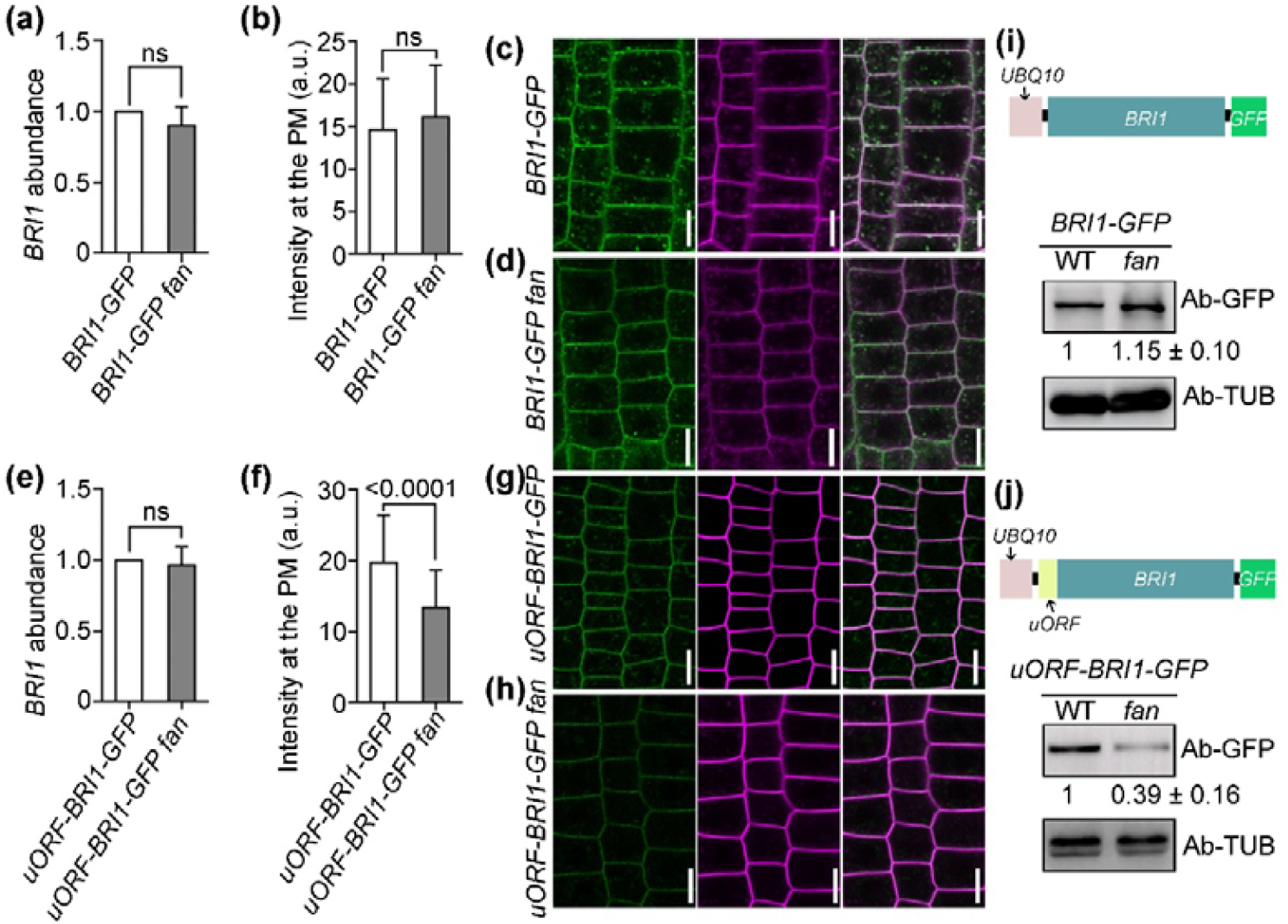
*FAN* is important in maintaining *BRI1* translational efficiency. (a) Relative *BRI1-GFP* abundance between *UBQ10:BRI1-GFP* and *UBQ10:BRI1-GFP fan*. (b) GFP intensity at the plasma membrane (PM) of *UBQ10:BRI1-GFP* or *UBQ10:BRI1-GFP fan*. (c-d) Representative CLSM images of *UBQ10:BRI1-GFP* (c) or *UBQ10:BRI1-GFP fan* (d) roots. (e) Relative *BRI1-GFP* abundance between *UBQ10:uORF-BRI1-GFP* and *UBQ10:uORF-BRI1-GFP fan*. (f) GFP intensity at the PM of *UBQ10:uORF-BRI1-GFP* or *UBQ10:uORF-BRI1-GFP fan*. (g-h) Representative CLSM images of *UBQ10:uORF-BRI1-GFP* (g) or *UBQ10:uORF-BRI1-GFP fan* (h) roots. For (a, e), RNAs were extracted from seedlings at 7 DAG; results are means ± SE (n = 3). For (b, f), results are means ± SD (n = 54 to 100 cells). For (a, b, e, f), Students’ *t*-test was performed (P > 0.05 indicated by ns). For (c-d, g-h), roots were pulse-labeled with FM4-64 before imaging; images shown from left to right are: the GFP channel, the RFP channel, and merges of the GFP/RFP channels. (i-j) Western blot assays showing the relative level of BRI1-GFP in *UBQ10:BRI1-GFP* transgenic seedlings (i) or in *UBQ10:uORF-BRI1-GFP* transgenic seedlings (j). A cartoon illustrating the expression cassette for *UBQ10:BRI1-GFP* (i) or *UBQ10:uORF-BRI1-GFP* (j) is shown on top of the Western blot assays. Tubulin (TUB) was used as the internal control. BRI1-GFP levels in wild-type backgrouns were set as 1. Results shown are means ± SE (n = 3). Bars = 10 µm.

By CLSM imaging of roots, we determined that BRI1-GFP signals were comparable between *UBQ10:BRI1-GFP* seedlings and *UBQ10:BRI1-GFP fan* seedlings (Fig. 7b, 7c, 7d). The comparable abundance of BRI1-GFP in *UBQ10:BRI1-GFP* seedlings and *UBQ10:BRI1-GFP fan* seedlings was further supported by biochemical assays using anti-GFP antibodies (Fig. 7i). By contrast, BRI1-GFP abundance was significantly reduced in *UBQ10:uORF-BRI1-GFP fan* seedlings than in *UBQ10:uORF-BRI1-GFP* seedlings either by CLSM imaging (Fig. 7f, 7g, 7h) or by biochemical assays (Fig. 7j), despite that the transcript abundance of *BRI1-GFP* was comparable between wild type and *fan* (Fig. 7e). These results demonstrate that FAN-FIP1, by mediating ribosomal biogenesis, are important for maintaining translational efficiency of *BRI1* and thus BR signaling.

### BL promotes pre-rRNA processing in a BRI1-dependent way

BRs were reported to mediate post-transcriptional accumulation of chloroplast rRNAs (Komatsu et al., 2010). However, it was unclear whether BRs influenced nuclear rRNA processing. To test this possibility, we applied BL and examined pre-rRNA processing by quantitative reverse-transcription PCRs. Treatment with 100 nM BL for 2 hrs increased the total rRNA levels without affecting the transcription of 45S pre-rDNA or the levels of pre-rRNAs (Fig. 8a-b), indicating that BL promotes ribosomal biogenesis. Because BRI1 is crucial for BR signaling (Li & Chory, 1997, Wang et al., 2001, Wang et al., 2006), we examined whether the effect of BL on pre-rRNA processing relied on BRI1 by using the *BRI1* null mutant *bri1-116* (Wang et al., 2001, Oh et al., 2012, Li et al., 2020). Indeed, BL-induced increase of pre-rRNA processing was completely abolished in *bri1-116* (Fig. 8b), suggesting that BL promoted pre-rRNA processing in a BRI1-dependent way.

**Figure 8.**
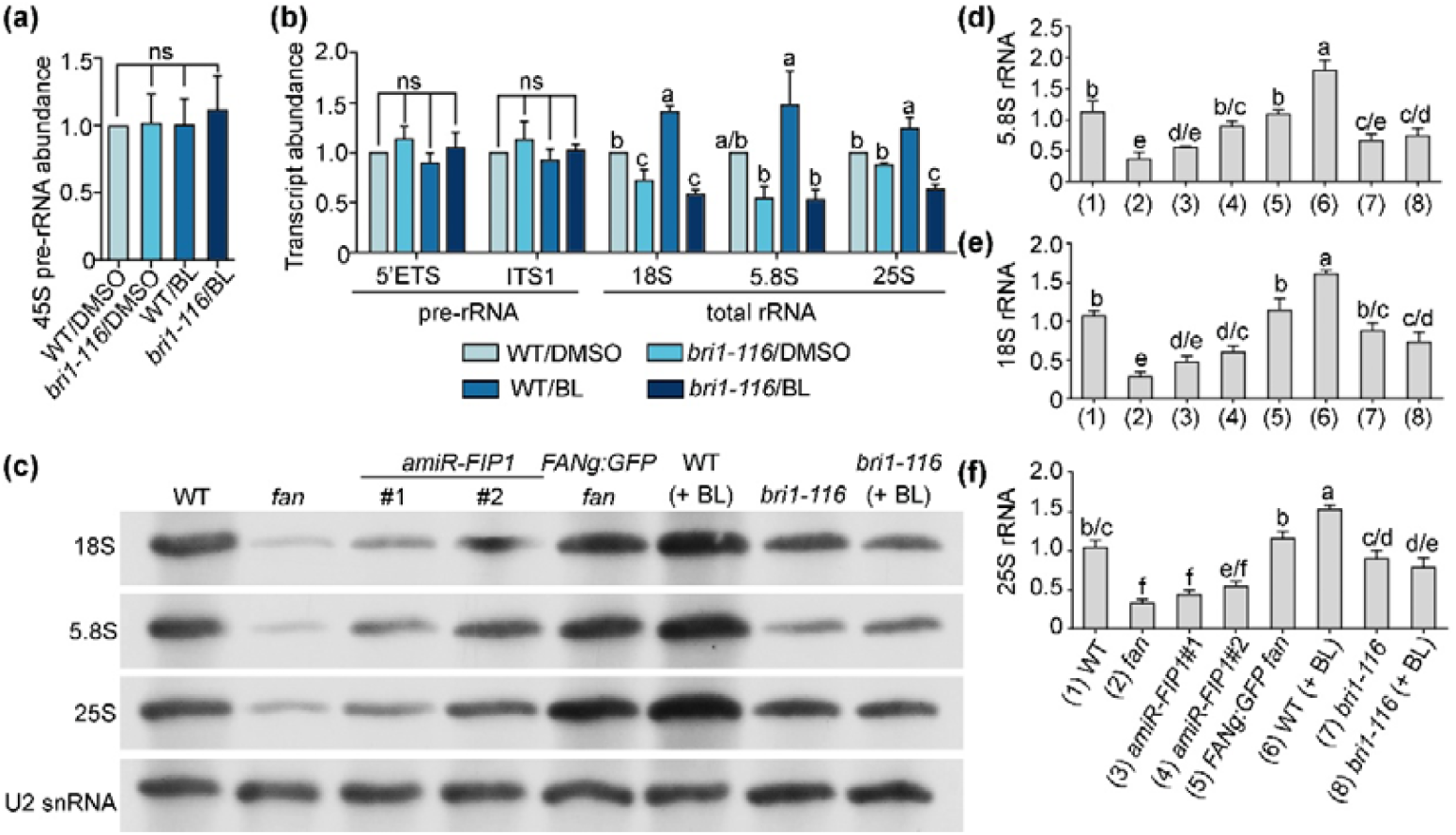
BL promotes pre-rRNA processing in a BRI1-dependent way. (a-b) RT-qPCRs of 45S pre-rRNA (a) or rRNA fragments (b) abundance in wild type or *bri1-116* seedlings at 7 d after germination (DAG) treated with DMSO or BL. Expression levels are related to that of *EF-1α* (left) or 45S pre-rRNA (right). Results shown are means ± SD (n = 3). Three biological replicates were examined with similar results. Different letters indicate significant different groups (OneWay ANOVA, Tukey’s multiple comparisons test, P < 0.05); ns indicates no significant difference among different genotypes (OneWay ANOVA, Tukey’s multiple comparisons test, P > 0.05). For chemical treatment, sterilized seeds were grown on 1/2 MS plates for 7 days, followed by a 2-hour treatment with DMSO or 100 nM BL before RNA extraction. (c) A representative Northern blot for mature rRNAs. Total RNAs were extracted from 7 DAG seedlings. RNAs were separated in 1% agarose polyacrylamide gels and hybridized with probes specific for particular regions of 18S rRNA, 5.8S rRNA, and 25S rRNA. Hybridization for U2 snoRNA was used as the loading control. (d-f) Abundance of 18S rRNA (d), 5.8S rRNA (e) or 25S rRNA (f). Results shown are means ± SD (n = 3 biological replicas). Different letters indicate significantly different groups (OneWay ANOVA, Tukey’s multiple comparisons test, P < 0.05).

To further verify the compromised rRNA processing in *bri1-116* as well as the enhanced processing of rRNAs by exogenous BL treatment, we applied Northern blot assays to examine the level of mature 18S, 5.8S, and 25S rRNAs (Fig. 8c), in which U2 snRNA was used as a loading control (Maekawa et al., 2024, Wang et al., 2024, Zakrzewska-Placzek et al., 2025). We also included *fan, amiR-FIP1* transgenic lines, and *FANg:GFP fan*, to corroborate the results (Fig. 8c). Consistent with the RT-qPCR results, exogenous application of BL induced a significant increase of 18S, 5.8S, and 25S rRNAs (Fig. 8d-f) whereas the BR signaling deficient mutant *bri1-116* not only showed a significantly reduced accumulation of mature rRNAs but did not respond to exogenous BL (Fig. 8d-f). The accumulation of mature rRNAs in *fan* and *amiR-FIP1* lines was significantly reduced compared to that of wild type or *FANg:GFP fan* plants (Fig. 8d-f), demonstrating the fidelity of the experiments.

### FAN-FIP1 participate in heat tolerance

BR signaling not only mediates plant development but also promotes heat tolerance (Chen et al., 2022). In addition, heat stresses inhibit the efficient cleavage of the 5’ETS primary cleavage site in 45S pre-rRNA, leading to the accumulation of 45S pre-rRNA and interfering with the processing of rRNAs in Arabidopsis (Shanmugam et al., 2021, Darriere et al., 2022, Zakrzewska-Placzek et al., 2023, Zakrzewska-Placzek et al., 2025). Thus, we wondered whether the compromised ribosome biogenesis and BR signaling in *fan* and *amiR-FIP1* would lead to an increased sensitivity to heat.

To test this idea, we examined heat tolerance of *fan* and *amiR-FIP1* as compared to wild type. Indeed, compared to around 20% survival rate of wild-type seedlings heat-stressed for 8 hrs, less than 5% *fan* or *amiR-FIP1* seedlings survived after 6 days of recovery (Fig. 9a-b), indicating that FAN-FIP1 participated in heat tolerance.

**Figure 9.**
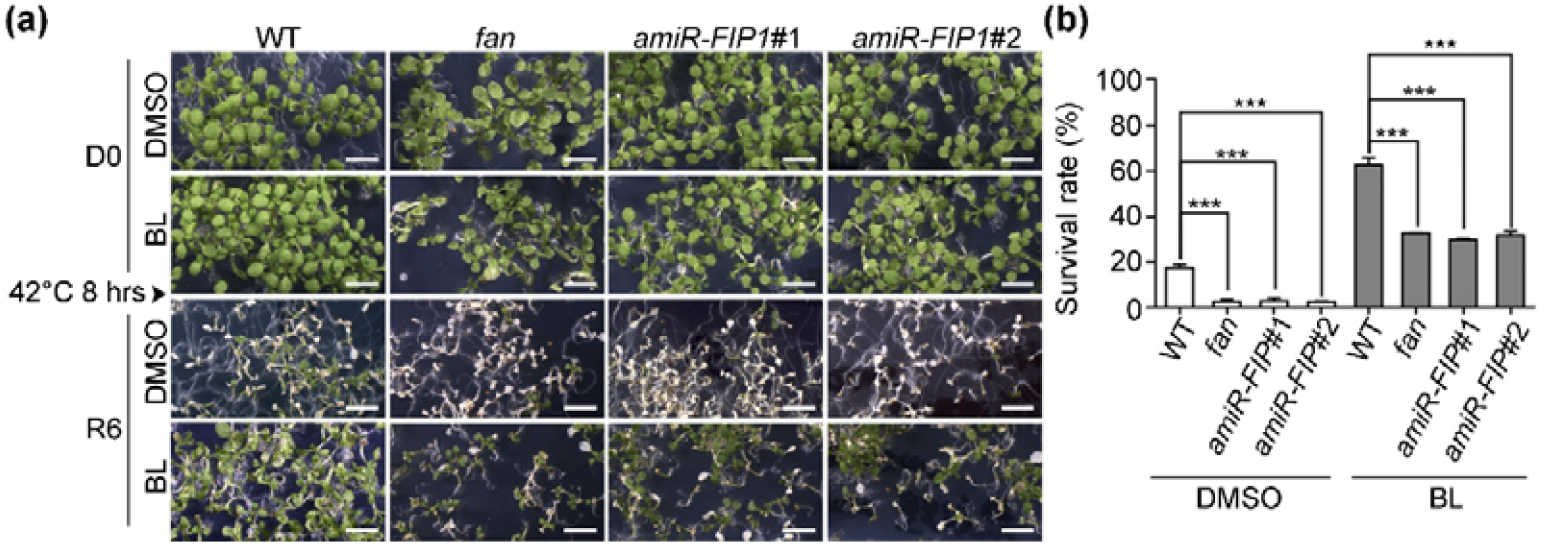
*FAN* loss-of-function and *FIP1* knock-down result in hypersensitivity to heat stresses. (a) A representative heat stress assay. Seedlings at 7 DAG (D0) under regular growth condition (22°C) on 1/2 MS supplemented with DMSO or 100 nM BL were placed at 42°C for 8 hrs and then transferred to 22°C for 6 days for recovery (R6). (b) Quantification of survival rate. Results are means ± SD (n = 3). More than 300 seedlings for each genotype were examined. Asterisks indicate significant difference (*t*-test, P < 0.0001). Bars = 5 mm.

Consistent with the previous report (Chen et al., 2022), wild-type seedlings treated with 100 nM BL significantly improved the survival rate under heat stress such that over 60% wild-type seedlings survived 42°C for 8 hrs at the presence of BL in contrast to around 20% with DMSO (Fig. 9a-b). Although the survival rate of *fan* or *amiR-FIP1* seedlings under heat stresses was also raised by BL treatment, the survival rate could hardly reach to 35% (Fig. 9a-b), further supporting the reduced heat tolerance of *fan* or *amiR-FIP1* to BL.

### FIP1 does not mediate DNA damage response

FAN was reported to protect genome integrity by mediating DNA damage response (DDR) through the ATR pathway (Liu et al., 2023), although the molecular components involved in FAN-mediated DDR were unknown. Since AATF proteins were also important for genome integrity in metazoans (Passananti & Fanciulli, 2007), we wondered whether FIP1 was an evolutionarily conserved partner for the role of FAN in DDR. To test this idea, we examined DDR of *amiR-FIP1* seedlings by using hydroxyurea (HU) treatment. HU is a genotoxic agent inducing cell death in Arabidopsis roots at 2 mM (Liu et al., 2023). Without HU treatment, *fan* roots showed a significantly higher cell death than those of wild type (Fig. S8). In comparison, knocking down *FIP1* did not cause an enhanced cell death (Fig. S8). HU treatment resulted in a significant increase of cell death in *fan* compared to that in wild type (Fig. S8). By contrast, knocking down *FIP1* showed a similar increase in cell death to that of wild type (Fig. S8). These results suggest that unlike FAN, FIP1 does not participate in DNA damage response.

## Discussion

Our study establishes that the Arabidopsis FAN-FIP1 complex, orthologous to yeast Bfr2-Lcp5 and metazoan AATF-NGDN, constitutes a core ribosome biogenesis factor (RBF) required for pre-rRNA processing and ribosomal subunit assembly (Fig. 1, Fig. 5). Studies in metazoans and in yeast demonstrate a direct role of FAN-FIP1 homologs in pre-rRNA processing as well as in ribosomal biogenesis, indicating an evolutionary conservation.

While this conserved role aligns with findings in other eukaryotes (Bammert et al., 2016, Kaiser et al., 2020), our work reveals novel plant-specific regulatory dimensions. First, FAN-FIP1 intersects with brassinosteroid (BR) signaling through uORF-mediated translational control of *BRI1* (Fig. 7), which has been demonstrated to be critical for BR signaling (Zhang et al., 2018, Zhang et al., 2020). Second, BR signaling reciprocally enhances pre-rRNA processing (Fig. 8). Third, this module contributes to heat tolerance (Fig. 9), a pathway not yet characterized in other systems. These discoveries position FAN-FIP1 as a critical integrator of ribosome biogenesis, hormone signaling, and stress adaptation in plants.

A key advance of this study is the elucidation of a reciprocal regulatory loop between FAN-FIP1 and BR signaling. We demonstrate that impaired ribosome biogenesis in *fan* or in *amiR-FIP1* transgenic plants reduces BRI1 receptor abundance via inefficient translation of its uORF-containing transcript (Fig. 7), leading to diminished BR sensitivity (Fig. 6). Crucially, we show that BR signaling promotes pre-rRNA processing in a BRI1-dependent manner (Fig. 8). This bidirectional crosstalk represents a plant-adaptive mechanism linking translational capacity to growth regulation. BR-driven growth demands increased ribosome production while FAN-FIP1 ensures efficient translation of BR signaling components. Although AATF orthologs influence translation in metazoans (Bammert et al., 2016, Kaiser et al., 2019), their connection to hormone signaling is uncharacterized. Thus, the FAN-FIP1-BR axis defines a unique plant pathway coordinating ribosome biogenesis with developmental plasticity. Despite that, key issues remain. It is highly likely that a lot more uORF-containing genes beyond *BRI1* would be affected in *fan* or *amiR-FIP1*. Ribosome profiling of mutants could identify selective translational targets. In addition, although BRI1 is required for BR-enhanced pre-rRNA processing (Fig. 8), the direct components in BR signaling for the regulation of pre-rRNA-processing are unclear. Key transcription factors in BR signaling are candidates whose activities would lead to enhanced expression of pre-rRNA-processing genes.

The FAN-FIP1 complex underpins both developmental and stress-response programs. Impaired ribosome biogenesis in *amiR-FIP1* causes developmental retardation (Fig. S6) and disrupts RAM maintenance (Fig. 4), resembling mutants with disrupted BR signaling (Tang et al., 2008, González-García et al., 2011, Hacham et al., 2011, Ryu & Hwang, 2013). Notably, FAN-FIP1 also mediates heat tolerance (Fig. 9), likely through its dual roles in maintaining ribosomal output and BR sensitivity. Heat stress disrupts pre-rRNA processing in plants (Darriere et al., 2022) and BRs promote thermotolerance by preserving translation (Chen et al., 2022). Our finding that FAN-FIP1 mutants show hypersensitivity to heat, which is partially rescued by BL yet never reaching wild-type resilience (Fig. 9), suggests that efficient ribosome biogenesis is a prerequisite for BR-mediated stress protection. This integrates two previously disconnected paradigms: ribosome assembly as a stress vulnerability point, and BRs as thermotolerance modulators. It has to be noted that whether heat sensitivity of *fan* or *amiR-FIP1* is solely attributable to ribosomal defects or exacerbated by disrupted BR responses remain to be determined.

The role of FAN in DNA damage response (DDR) (Liu et al., 2023) appears mechanistically distinct from its ribosome biogenesis function. While FAN interacts with FIP1 via its AATF domain to regulate rRNA processing (Fig. 2, Fig. S5), we now demonstrate that FAN also post-translationally stabilizes FIP1 such that GFP-FIP1 protein abundance is significantly reduced in *fan* despite unchanged transcript levels (Fig. 3). This regulatory layer ensures coordinated activity of the FAN-FIP1 complex and explains the shared ribosome biogenesis defects in both mutants. While FAN interacts with FIP1 via its AATF domain to regulate pre-rRNA processing (Fig. 2, Fig. S5), FIP1 is dispensable for DDR (Fig. S8). This implies that FAN employs distinct molecular partners for its nuclear functions, i.e. FIP1 for ribosomal maturation and unidentified ATR pathway components for genome maintenance. Whether these pathways converge developmentally remains an open question. FAN may act as a multifunctional scaffold, with its AATF domain dedicated to ribosome biogenesis and other domains (Fig. 2) mediating DDR, a model reconciling conservation with lineage-specific adaptations.

In conclusion, although FAN-FIP1’s core role in ribosome biogenesis is evolutionarily conserved, its integration with BR signaling and heat tolerance represents a plant-specific innovation. By demonstrating that (1) FAN-FIP1 enables BR signaling via uORF-dependent *BRI1* translation, and (2) BRs reciprocally boost rRNA processing, we unveil a feedback loop optimizing ribosome production for growth and stress adaptation. Future studies should dissect the molecular interfaces between BR signaling, ribosomal biogenesis, and DDR—pathways unified through FAN’s modular functions. These insights position ribosome biogenesis not merely as a housekeeping process, but as a dynamic hub for hormone-environment integration in plants.

### Experimental procedures

#### Plant growth and transformation

Arabidopsis (*Arabidopsis thaliana*) accession Columbia-0 (Col-0) was used as the wild type for all experiments. Plant materials, including *fan* and *FANg*:GFP, were described (Liu et al., 2023). Seedlings and plants were grown in a growth chamber at 20 ± 2ºC under long-day condition (16 h light/8 h dark). Plant transformation was as described (Zhou et al., 2013a). Stable transgenic plants were selected on half-strength MS medium supplemented with 30 μg/ml Basta salts (Sigma-Aldrich).

#### DNA manipulation

All constructs were generated using the Gateway technology (Invitrogen) unless otherwise specified. Entry clones were generated in the pENTR/D/TOPO vector (Invitrogen). The promoter for *FIP1* (*proFIP1*) was cloned with the primer pair P386/P387 from Columbia-0 genomic DNA, containing a 539 bp sequence upstream of its start codon. The *proFIP1* was introduced into the destination vector *GW:GUS* (Zhou et al., 2013a) to generated *proFIP1*:GUS. The full-length coding sequences (CDS) of *FAN and FIP1* were cloned by using the primer pair P306/P307 and P196/P197, respectively. The truncated *FAN* sequences encoding N-terminal 1-813 bp, the AATF domain (423-813 bp), and the TRAUB domain (1050-1275 bp) were cloned by using the primer pair P306/P1956, P1955/P1956, and P904/WYN-84, respectively. Sas10 domain (70-312 bp) of *FIP1* was cloned by using the primer pair P1958/P1959. The CDS of *BRI1* was cloned by using the primer pair ZP765/ZP766, the sequences of uORF-*BRI1* was cloned with the primer pair wyn-354/wyn-408. The CDS of *ARF3* was described previously (Xiong et al., 2020). Entry vectors were used in LR reactions with the destination vectors *UBQ10:GW-GFP, UBQ10:GFP-GW*, or *35S:GW-mRFP* (Karimi et al., 2002, Zhang et al., 2015) to generate *UBQ10:BRI1-GFP, UBQ10:uORF-BRI1-GFP, UBQ10:GFP-FIP1*, or *35S:ARF3-mRFP*. Constructs used in translational capacity assays, including *35S:ARF3-mRFP, 35S:uORF-ARF3-GFP* were described previously (Xiong et al., 2020). Two expression cassettes *35S:ARF3-mRFP* and *35S:uORF-ARF3-GFP* were stacked to generate the expression vector *35S:uORF-ARF3-GFP;35S:ARF3-mRFP*.

For vectors used in protein-protein interaction assays, including Y2H and BiFC, entry vectors were combined with the destination vectors *pDEST-GBKT7* or *pDEST-GADT7* (Clontech) for Y2H, or with the destination vectors *pSITE-cEYFP-C1* or *pSITE-nEYFP-C1* (Martin et al., 2009) for BiFC in LR reactions to generate corresponding expression vectors. To generate expression vectors used in FRET assays, the entry vector of *FAN* was used in LR reaction with the destination vector *35S:GFP-GW* to generate the expression vector *35S:GFP-FAN* whereas the entry vector of *FIP1* was used in LR reaction with *35S:mCherry-GW* to generate *35S:mCherry-FIP1*.

To generate *UBQ10:amiR-FIP1*, the *amiR-FIP1* fragments were amplified with the primers ZP6117/ZP6118/P1750-P1751 designated according to the online tool WMD3 (http://wmd3.weigelworld.org/cgibin/webapp.cgi). The resultant PCR fragments were inserted in pRS300 (Ossowski et al., 2008) and the *amiR-FIP1* fragments were amplified from pRS300 with the primer pair P350/P351. The resultant PCR fragments were inserted in pENTR/SD/D-TOPO to generate the entry vector, which was used in an LR reaction with the destination vector *UBQ10:GW* to generate *UBQ10:amiR-FIP1*.

All constructs were sequenced and analyzed using Vector NTI. All PCR amplifications were performed with Phusion hot-start high-fidelity DNA polymerase (Thermo Fisher Scientific) with the annealing temperature and extension times recommended by the manufacturer. All primers are listed in the Table S1.

#### RNA extraction and reverse transcription quantitative PCRs (RT-qPCRs)

For reverse transcription quantitative PCRs (RT-qPCRs) of *FIP1*, total RNAs were isolated from various tissues of wild type, include seedlings and roots at 7 days after germination (DAG), leaves at 14 DAG, stems at 25 DAG, inflorescences at 4-5 days after flowering, mature ovules, and siliques. For RT-qPCRs analyzing *FIP1* transcription levels in *amiR-FIP1* lines, total RNAs were isolated from roots of 7 DAG seedlings. For RT-qPCRs analyzing pre-rRNA, total rRNAs, *BRI1*, and BR marker genes in wild type versus *fan*, DMSO-versus BL-treatment assays, total RNAs were isolated from 7 DAG seedlings.

First-strand cDNA synthesis was performed with oligo(dT) and SuperScript III reverse transcriptase with on-column DNase digestion (Invitrogen). RT-qPCRs were performed by using a Bio-Rad CFX96 real-time system with SYBR Green real-time 13 PCR master mix (Toyobo) as described (Zhou et al., 2013a). Primers used in RT-qPCRs are the following: P395/P396 for *FIP1*, P937/P938 for 45S pre-rRNA, P1090/P1091 for *5’ETS1*, P941/942 for *ITS1*, P945/946 for *18S rRNA*, P1088/P1089 for *5*.*8S rRNA*, P5165/5166 for *25S rRNA*, WYN-427/WYN-428 for *BR6ox1*, WYN-429/WYN-430 for *ROT3*, WYN-406/WYN-407 for *BRI1*, NKP240/NKP241 for *SAUR-AC1. GAPDH* was used as the internal reference gene for RT-qPCRs analyzing the expression of *FIP1, BR6ox1, ROT3, BRI1 and SAUR-AC1. EF-1α* was used as reference gene for RT-qPCRs analyzing pre-rRNA and total rRNA. Analyses of expression were performed using Prism 7 (GraphPad Software). All primers are listed in the Table S1.

#### GUS histochemistry

Plant materials including mature ovules, embryos, inflorescences, seedlings, roots, and leaves were incubated for 4 hr at room temperature in the dark with 5-bromo-4-chloro-3-indolyl-glucronide (X-Gluc) before imaging. The final working concentration of X-Gluc was 1 mg/ml (Wang et al., 2013, Zhou et al., 2013a). Images were captured with an Olympus BX53 microscope or an Olympus SZX16 microscope as described (Wang et al., 2013, Zhou et al., 2013a).

#### Polysome profiling

Polysome profiling was performed as described (Li et al., 2018, Xiong et al., 2020), with slight modifications. Briefly, seedlings from 7 DAG wild-type, *fan, FANg:GFP fan*, or *amiR-FIP1* plants were ground in liquid nitrogen, followed by resuspension in polysome extraction buffer (0.2 M Tris-HCl, pH 7.5, 50mM KCl, 25 mM MgCl_2_, 50mM EGTA, pH 8.0, 0.05 mg/mL cycloheximide, 0.025 mg/mL chloramphenicol, 0.5% NP40 [v/v]). The extract was loaded onto a 10% - 50% sucrose gradient and spun in a Beckman Optima XPN-100 Ultracentrifuge (SW41Ti rotor) at 38,000 rpm for 3 hr at 4°C. The gradients were fractionated using a density gradient fractionation system with continuous monitoring of absorbance at 260 nm.

#### Flow cytometry for cell ploidy

Seedlings were chopped with a razor blade in 100 µL of nuclei extraction buffer (45 mM MgCl_2_, 30 mM sodium citrate, 20 mM MOPS pH 7.0 and 0.2% Triton X-100) on ice and resuspended with 500 µL of nuclei extraction buffer (Galbraith, 1990). All tissues and solution were then transferred into a 1.5 mL tube and mixed well by using vortex. The supernatant was filtered over a 33 µm mesh, and incubated with DAPI at a final concentration of 0.001 mg/mL at 4 °C for 5-10 min. The nuclei were analyzed with flow cytometer (BD FACS AriaIII). At least 10000 nuclei were scored in triplicates for each sample.

#### Phenotypic analyses and pharmacological treatment

To measure primary root length, roots from 7 DAG seedlings were imaged and measured by using Image J (http://rsbweb.nih.gov/ij/). For RAM analyses, roots from 5 DAG seedlings were staining by PI (10 mg/mL from Sigma-Aldrich, dissolved in water) for 5 min, then excited at 561 nm. For cell numbers and cell length within RAM, cells from quiescent center (QC) to the first elongated cell in the cortical cell layer were measured. For each genotype, 15 to 20 roots were used for measurement.

For nucleolus staining with acridine orange (AO, Macklin, CAS:10127-02-3), 5 DAG seedlings were incubated with AO a final concentration of 0.5 μM/mL for 20 min at room temperature, washed three times with distilled water, incubated with DAPI (1 mg/mL) for 15 min at room temperature in the dark, and then imaged with CLSM. The excitation and emission wavelengths settings are 458 nm/600 to 680 nm for AO signals and 405 nm/420 to 480 nm for DAPI signals.

For BL treatment on hypocotyl elongation, root growth, and RAM development, BL was prepared in DMSO as a 100 mM stock while the working concentrations were 100 nM for hypocotyl elongation and root growth assays, 0.8 nM for RAM development assays. DMSO was equally diluted as the controls. Arabidopsis seeds were placed on a 1/2 MS medium with DMSO, 100 nM BL, or 0.8 nM BL and kept at 4°C in darkness for 1 to 2 days to synchronize germination. For hypocotyl elongation assay, plates were transferred to a growth chamber with a 8-h light followed with 5 d in the dark before imaging. For root growth and RAM development assays, plates were transferred to a growth chamber with a 16-h light/8-h dark cycle at 22°C for 7 d or 5 d, respectively.

For HU treatment, HU was prepared in water as a 650 mM stock while the working concentration was 2 mM. Arabidopsis seeds were placed on a 1/2 MS medium and kept at 4°C in darkness for 1 to 2 d before being transferred to a growth chamber with a 16-h light/8-h dark cycle at 22°C for 4 d. Then seedlings were transferred to 1/2 MS medium with 2 mM HU for 24 hours before imaging. In order to quantify cell death area, roots were stained with 10 μM PI before CLSM imaging.

#### Heat tolerance assays

For seedling survival assays under heat stress, seedlings at 7 DAG on 1/2 MS medium containing DMSO or 100 nM BL were transferred to 42°C for 8 hrs in the dark and then transferred to a growth chamber with a 16-h light/8-h dark cycle at 22°C for 6 days for recovery. To assess survival rate, the number of fully etiolated, partially etiolated, and non-etiolated seedlings were measured. The survival rate was calculated as (the number of non-etiolated seedlings + 1/2 number of partially etiolated seedlings)/total number of seedlings.

#### Protoplast transformation

Arabidopsis protoplasts were prepared according to a previous report (Wang et al., 2017) with slight modifications. Briefly, rosette leaves from 3 weeks after germination (WAG) of Arabidopsis plants were cleaved and placed into 10 mL of the enzymatic solution (1% cellulase “Onozuka” R10 [Yakult], 0.25% macerozyme “Onozuka” R10 [Yakult], 0.4 M mannitol, 10 mM CaCl_2_, 20 mM KCl, 0.1% BSA, and 20 mM MES, pH 5.7) for 1-2 hr to obtain protoplasts. The protoplasts were then collected by centrifugation at 100 g for 2 min, washed once with 5 mL cold W5 solution (154 mM NaCl, 125 mM CaCl_2_, 5 mM KCl, 5 mM glucose, and 2 mM MES, pH 5.7), re-suspended with 5 ml cold W5 and incubated on ice at least 30 min. The W5 solution was sucked out and then the protoplasts were re-suspended in MMg solution (0.4 M mannitol, 15 mM MgCl_2_, and 4 mM MES, pH 5.7) to a final concentration of 2 - 5 × 10^5^ cells/mL. Transformation of Arabidopsis protoplasts were performed as described (Yoo et al., 2007). Plasmids containing designated expression cassettes were introduced into protoplasts by PEG-mediated transformation. Transformed protoplasts were cultured for 12 - 16 hr in a 22°C incubator and then examined.

#### Protein interaction assays

Y2H assays were performed as previously described (Park et al., 2014) with slight modifications. Briefly, different combinations of bait and prey vectors were co-transformed into the Y2H Gold yeast strain (Clontech). Protein-protein interactions were determined based on the growth of transformants after 3 d on YSD-WLHA.

For BIFC assays, a P19 protein was used to suppress gene silencing as described (Park et al., 2014). U1-70k-mCherry was used to label the nucleus as described (Wang et al., 2012). *pSITE-nEYFP-FAN/FAN*^N1-271^*/FAN*^N142-271^*/FAN*^350-425^ and *pSITE-cEYFP-FIP1/FIP1*^SAS10^ were infiltrated into tobacco leaves together with P19 and U1-70k-mCherry. Combinations of *pSITE-nEYFP-FAN/FAN* ^*N1-271*^ */FAN*^*N142-271*^ */FAN*^*350-425*^ and *pSITE-cEYFP-C1 or pSITE-cEYFP-FIP1/FIP1*^SAS10^ and *pSITE-nEYFP-C1* were used as negative controls. Confocal imaging was performed 48 hr after infiltration.

For FRET assays, the expression vectors *35S:GFP-FAN* (donor) and *35S:mCherry-FIP1* (acceptor) or *35S:mCherry* (acceptor) were transiently transformed in Arabidopsis protoplasts as described (Liu et al., 2021). Images in donor (excitation 488 nm; emission 500 to 550 nm), acceptor (excitation 561 nm; emission 580 to 620 nm), and FRET (excitation 488 nm; emission 580 to 620 nm) channels were captured. The energy transfer efficiency were calculated as follows: Efficiency=1-[e/(e+FRET)], FRET=*f*-DSBT-ASBT, ASBT=(*c/d*)×*g*, DSBT=(*b/a*)×*e*. Where *a* is the fluorescence intensity of the donor channel image excited at 488 nm when the donor is transformed alone, *b* is the fluorescence intensity of the acceptor channel image excited at 488 nm when the donor is transformed alone, *c* is the fluorescence intensity of the acceptor channel image excited at 488nm when the acceptor transformed alone, *d* is the fluorescence intensity of the acceptor channel image excited at 561 nm when the acceptor transformed alone, *e* is the fluorescence intensity of the donor channel image excited at 488 nm when the donor and acceptor are transformed together, *f* is the fluorescence intensity of the acceptor channel image excited at 488 nm when the donor and acceptor are transformed together, *g* is the fluorescence intensity of the acceptor channel image excited at 561 nm when the donor and acceptor are transformed together.

#### Protein biochemical assays

For protein abundance assays, Arabidopsis 7 DAG seedlings were grinded into powder, then added lysis buffer (10 mM HEPEs, pH=7.5, 100 mM NaCl, 1 mM EDTA, 10% Glycerol, 0.5% Triton X-100, and 1 × complete protease inhibitor cocktails) and mixed well. The supernatant was collected by centrifugation (12000 rpm) (Thermo Fisher, Sorvall Legend Micro 21R, Dual Row 18 × 2.0/0.5mL Rotor) at 4°C for 10 min. Supernatant were boiled with 1 × SDS loading buffer at 100°C for 10 min and detected by Western-Blot. Immunolabeling of BRI1-GFP/uORF-BRI1-GFP was performed using an anti-GFP antibody (Abmart, M2004, 1:5000 dilution). Anti-Tubulin (TransGen, HC101, 1:5000 dilution) were used as the loading control. Quantification of total proteins was performed using ImageJ.

#### Northern blot assays

Total RNAs were extracted from 7 DAG seedlings; 10 μg of total RNAs was taken and mixed with 3×formaldehyde loading buffer. The mixture was denatured at 65°C for 15 min and then placed on ice for 5 min before agarose gel electrophoresis. The agarose gel was placed on a nylon membrane (GE Healthcare) that had been pre-soaked in transfer buffer. The agarose gel was then transferred at 50-58 mbar for 2 hrs (1 mL transfer buffer was added every 10 min). After completion, the nylon membrane was washed with 1×MOPS buffer for 10 sec, dried with absorbent papers, and fixed by UV cross-linking (Thermo Fisher) at 120000 μJ/cm^2^. The membrane was placed in a hybridization tube and hybridized with ULTRAhyb hybridization solution (Ambion) (10 mL/100 cm^2^ membrane) at 42°C for 4 hrs. During this time, the probe was diluted 10 times with 10 mM EDTA, denatured at 90°C for 10 min, and placed on ice before use. The denatured probe was added to the hybridization solution and hybridized at 42°C overnight (16-20 hrs). After hybridization, the membrane was washed twice with Low Stringency Wash Solution (Ambion) at room temperature for 5 min each, and then washed twice with High Stringency Wash Solution (Ambion) at 42°C for 20 min each. The membrane was wrapped with cling film, placed on X-ray film (Fujifilm), and exposed at −80°C for 12-24 h. After development, the bands were observed. γ-^32^P 5’ end-labelled oligonucleotides (Table S1) were used as probes against mature ribosome RNAs and U2 snoRNA-specific probes. Quantification of Northern blots was performed using ImageJ software.

#### Fluorescence microscopy and quantification

All fluorescent images were captured using a Zeiss LSM 880 confocal laser scanning microscope (CLSM) with a 20 or 40/1.3 oil objective. Fluorescence of GFP, YFP, RFP, mCherry, or PI staining was captured using the following excitation/emission settings: 488 nm/505 to 550 nm for GFP, 514 nm/530 to 590 nm for YFP, 561 nm/575 to 650 nm for RFP, and 561 nm/600 to 650 nm for mCherry and PI staining. GFP-RFP double-labeled protoplasts were captured alternately using line-switching with the multi-track function. Image processing was performed with the Zeiss LSM image processing software (Zeiss). For quantification of green and red fluorescence intensity from WT or *fan* protoplasts, a region of interest (ROI) of the same size was defined in the nucleus (Nuc) within a protoplast. Ratio of fluorescence intensity between the RFP and GFP (RFP/GFP) was calculated using ImageJ. Up to 30 protoplasts for each genetic backgrounds were measured.

#### Phylogenetic analysis

Phylogenetic analysis was performed as described (Xiong et al., 2019, Liu et al., 2024a), with slight modifications. Phylogenetic analysis was performed using MEGA7 based on protein sequences of FIP1 homologs, calculated with the maximum likelihood method. Tree topology robustness was tested by bootstrap analysis of 1000 replicates. The protein sequences of FIP1 were obtained from TAIR, protein homologs of FIP1 from other species were identified by performing BLASTP searches against the NCBI protein database (http://www.ncbi.nlm.nih.gov/) using default parameters.

For protein-protein interaction prediction analysis, the amino acid sequences of the target proteins were downloaded from NCBI as FASTA files. AlphaFold3 (https://alphafoldsever.com) was used to predict protein structures. PyMOL was used to visualize the protein-protein interaction generated by AlphaFold3.

## Statistical analysis

Quantification data were analyzed using GraphPad Prism 6.02 (www.graphpad.com/scientificsoftware/prism/). Statistical analysis was performed with Student’s *t*-tests or OneWay ANOVA (Turkey’s multiple comparisons test).

## Accession numbers

Arabidopsis Genome Initiative locus identifiers for the genes mentioned in this article are as follows: At5g61330 for *FAN*, At1g07840 for *FIP1*, At2g33860 for *ARF3*, At4g37490 for *CycB1;1*, At1g08560 for *KNOLLE*, At5g38970 for *BR6ox1*, At4g36380 for *ROT3*, At3g04120 for *GAPDH*, At5g60390 for *EF-1α*, At4g39400 for *BRI1*, At4g38850 for *SAUR-AC1*.

## Author contributions

Y.Z., X.F., and Y.-N.W., designated this research; Y.-N.W. performed most of the experiments with the assistance of F.X., J.-Y.L., and S.L., Y.Z. and Y.-N.W. wrote the paper with input from all other authors.

## Acknowledgements

We thank Prof. Xugang Li for the *fan* and *FANg:GFP* seeds. This work is supported by National Natural Science Foundation of China (32400290 to F.X.), Taishan Scholar Project of Shandong Province of China (tsqn202408133), and by Shandong Provincial Natural Science Foundation (2025HWYQ-055 to F.X., ZR2024MC093 to S.L.). The authors declare no conflict of interest.

## Data Availability

All data generated or analysed during this study are included in this published article and its supplementary information files.

## Supplemental Information

Figure S1. Genomic *FAN* fully complements pre-rRNA processing defects in *fan*.

Figure S2. *fan* is compromised in cell cycle progression.

Figure S3. Phylogenetic analysis of FIP1.

Figure S4. *FIP1* is constitutively expressed.

Figure S5. FAN interacts with FIP1.

Figure S6. *FIP1* knock-down causes retarded growth.

Figure S7. *FIP1* knock-down results in enlarged nucleolus.

Figure S8. *FIP1* downregulation does not show enhanced DNA damage.

Table S1. Oligos used in this study.

