## Supplemental figures for "The nucleolar complex FAN-FIP1 mediates ribosome biogenesis in Arabidopsis and is critical for BR signaling and heat tolerance"

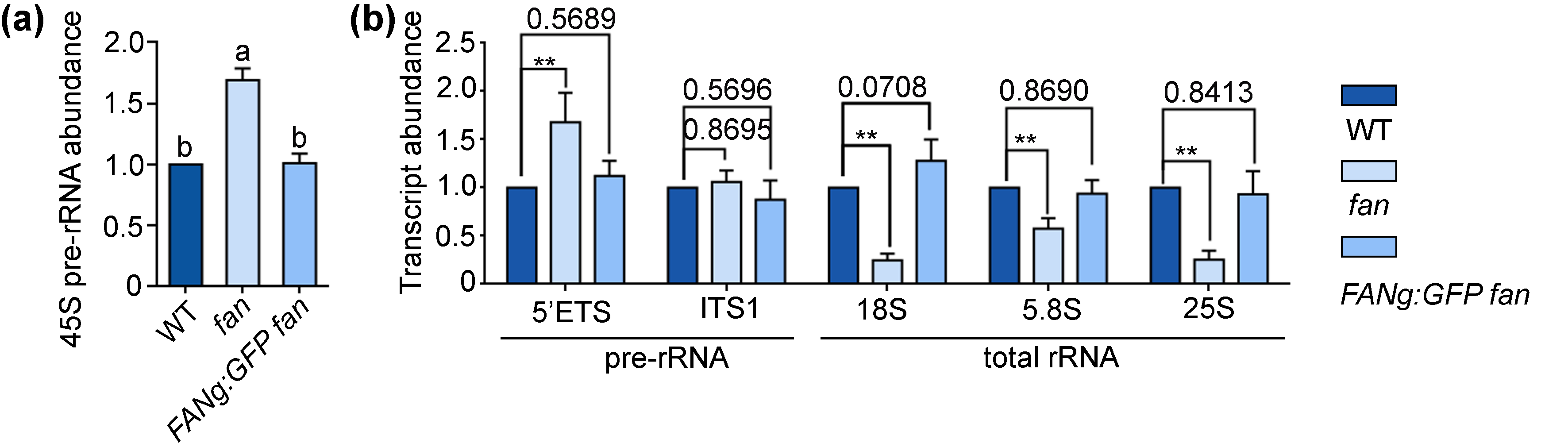

**Figure S1.** **Genomic *FAN* fully complements pre-rRNA processing defects in *fan*.**

(a-b) Reverse transcription quantitative PCRs (RT-qPCRs) of 45S pre-rRNA (a) or rRNA fragments (b) abundance in wild type, *fan*, or *FANg:GFP fan* seedlings at 7 days after germination (DAG). Expression levels are related to that of *EF-1α* (a) or 45S pre-rRNA (b). Results shown are means ± standard errors (SE, n = 3). *p* values above 0.01 are shown on top of the columns; *p* values < 0.01 are indicated by ** (unpaired *t*-test).

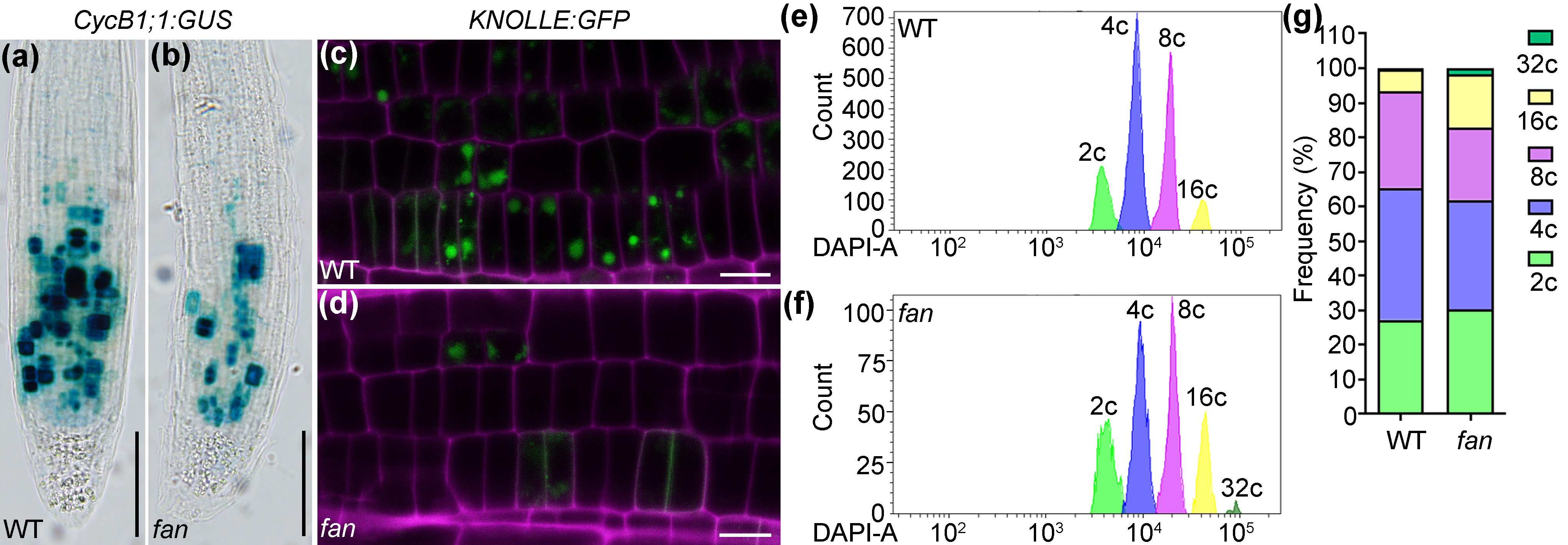

**Figure S2. *fan* is compromised in cell cycle progression**

(a-b) Representative histochemical GUS staining of *CycB1;1:GUS* (WT, a) or *CycB1;1:GUS* *fan* (*fan*, b) root tips from 7 days after germination (DAG) seedlings. (c-d) Representative confocal laser scanning microscopy (CLSM) images of *KNOLLE:GFP* (WT, c) or *KNOLLE:GFP fan* roots (*fan*, d) from 5 DAG seedlings. Images shown are merges of the GFP (green) and RFP (PI staining, magenta) channel images. (e-f) A one-dimensional frequency distribution of DNA content (linear scale) of nuclei from 7 DAG seedlings of wild type (e) or *fan* (f) by flow cytometric analyses*.* From the left, the five peaks of fluorescence correspond to nuclei having DNA contents of 2C, 4C, 8C, 16C, 32C, respectively. (g) Quantification of flow cytometry peaks 2C-32C of 7 DAG seedlings nuclei stained with DAPI. *fan* contains more 2C cells and less 4C and 8C cells. Bars = 50 µm (a-b); 10 µm (c-d).

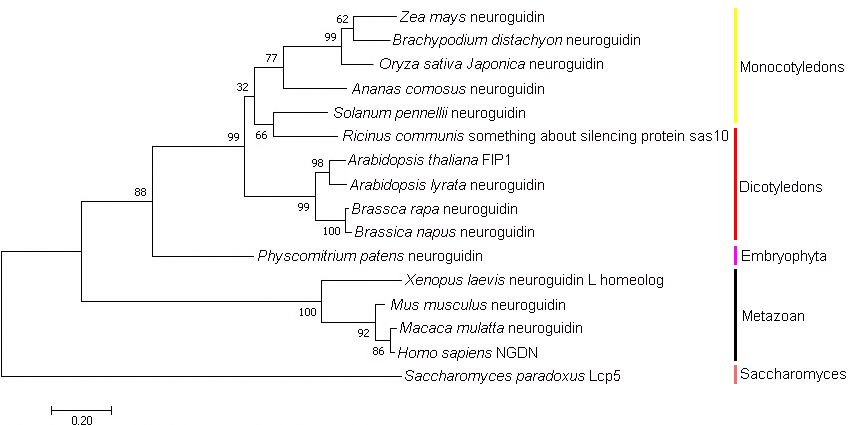

**Figure S3. Phylogenetic analysis of FIP1.**

Phylogenetic analysis of FIP1 orthologues using the maximum likelihood method with the MEGA7.0 software. All species protein sequences were obtained from the National Center for Biotechnology Information. These sequences including *Arabidopsis thaliana* (BAH19496.1), *Arabidopsis lyrata* (XP_002889668.1), *Zea mays* (PWZ11113.1), *Solanum pennellii* (XP_015071704.1), *Brachypodium distachyon* (XP_003567421.1), *Oryza sativa Japonica Group* (XP_015648588.1), *Ananas comosus* (OAY65329.1), *Ricinus communis* (EEF35039.1), *Brassica rapa* (XP_009148031.1), *Brassica napus* (XP_013712278.1), *Physcomitrium patens* (XP_024366029.1,), *Xenopus laevis* (NP_001088857.1), *Mus musculus* (NP_081166.1), *Macaca mulatta* (NP_001253183.1), *Homo sapiens* (NP_001036100.1), *Saccharomyces paradoxus* (XP_033765948.1). Tree topology robustness was tested by bootstrap analysis of 1,000 replicates. All parameters correspond to default definitions. The different colored vertical lines on the right represent the attributes of the species.

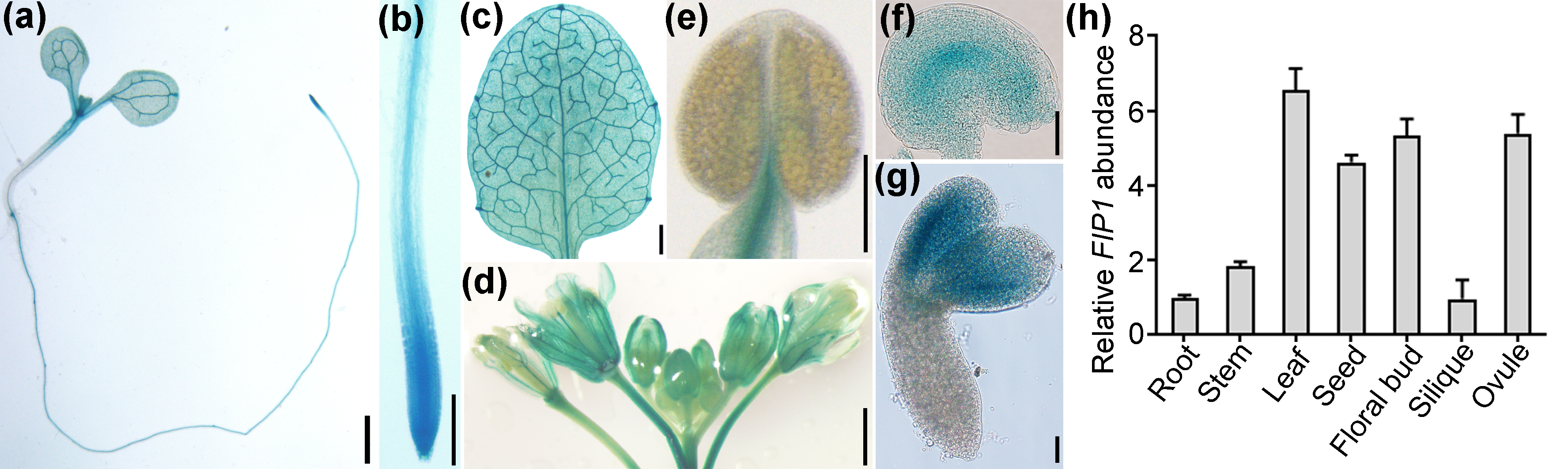

**Figure S4. *FIP1* is constitutively expressed.**

(a-g) Representative histochemical GUS staining of a seedling (a), a primary root tip (b), a rosette leaf (c), an inflorescence (d), a mature anther (e), a mature ovule (f), or a developing embryo (g) from the *proFIP1:GUS* plants. (h) Reverse transcription quantitative PCRs (RT-qPCRs) of *FIP1* among different tissues. Expression levels are related to that of *GAPDH*. Results shown are means ± standard errors (SE, n = 3). Each biological replicates were repeated three times with similar results. Bars = 1 mm (a, c, d), 200 µm (b, e), 20 µm (f, g).

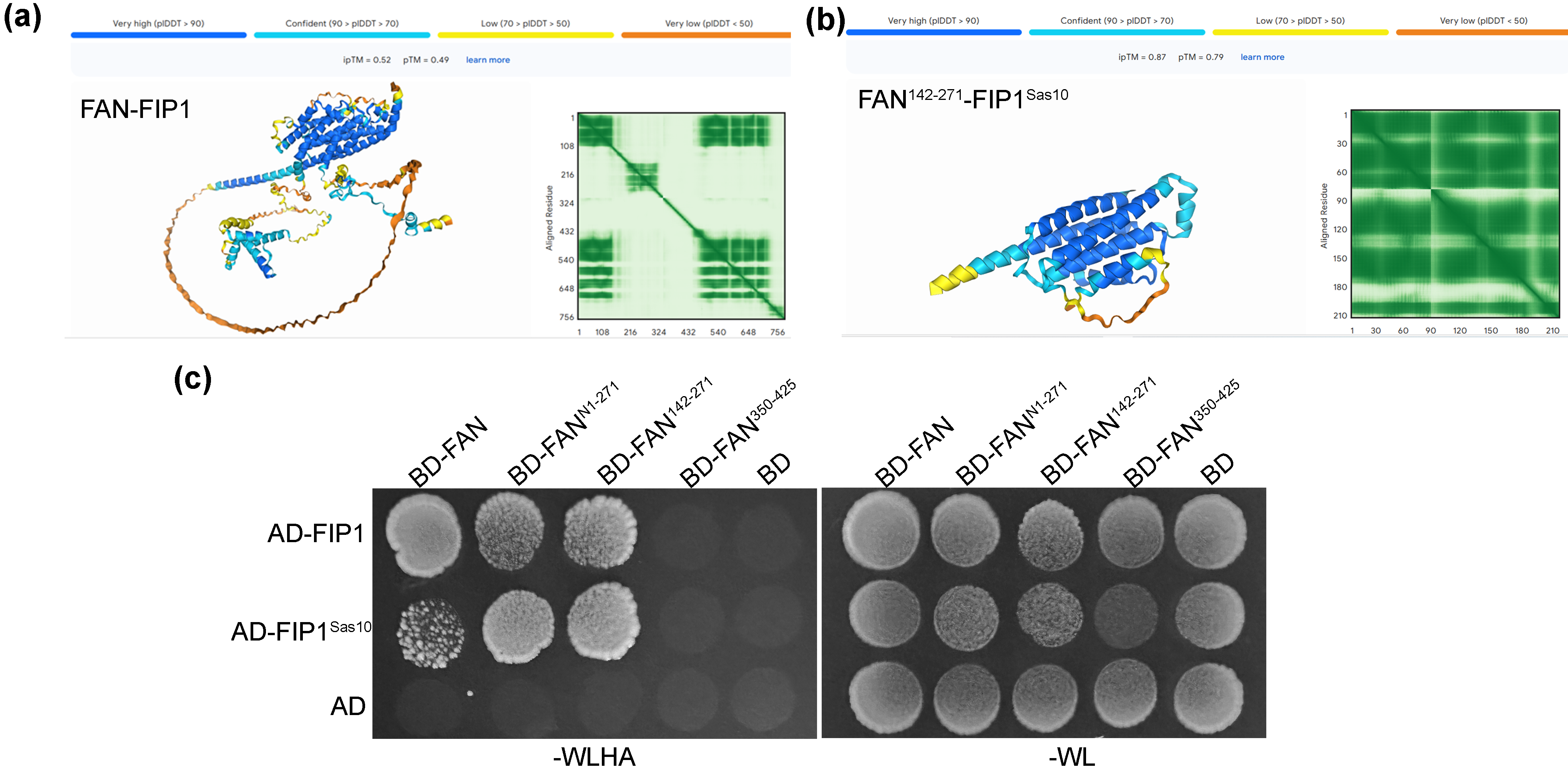

**Figure S5. FAN interacts with FIP1.**

(a-b) AlphaFold3.0 prediction of FAN-FIP1 (a) or FAN^142-271^-FIP1^Sas10^ interaction (b). (c) A representative yeast two hybrid (Y2H) assay. BD-FAN, BD-FAN^N1-271^, BD-FAN^142-271^ or BD-FAN^350-425^ was co-transformed with AD-fusions of FIP1 or FIP1^Sas10^. The co-transformed yeast strains were growing on selective (-WLHA) or non-selective (-WL) plates. -WLHA indicates YSD medium lacking Trp (W), Leu (L), His (H), and Ade (A), -WL indicates YSD medium lacking Trp and Leu. AD and BD were used as controls. Results are representative of three biological replicates.

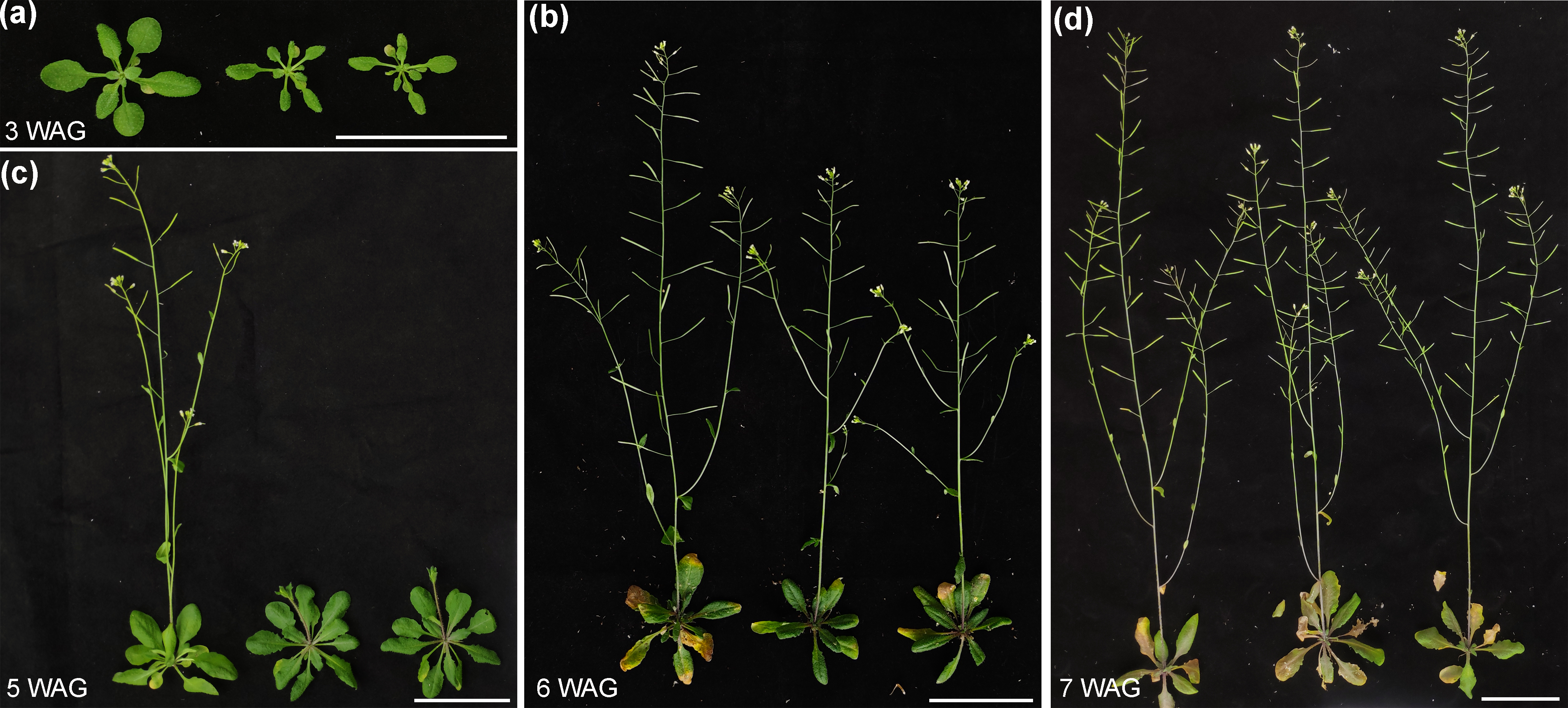

**Figure S6. *FIP1* knock-down causes retarded growth.**

(a-d) From left to right: a representative plant of wild type and two independent transgenic lines of *amiR-FIP1* at 3 weeks after germination (WAG) (a), 5 WAG (c), 6 WAG (b), or 7 WAG (d). Bars = 5 cm.

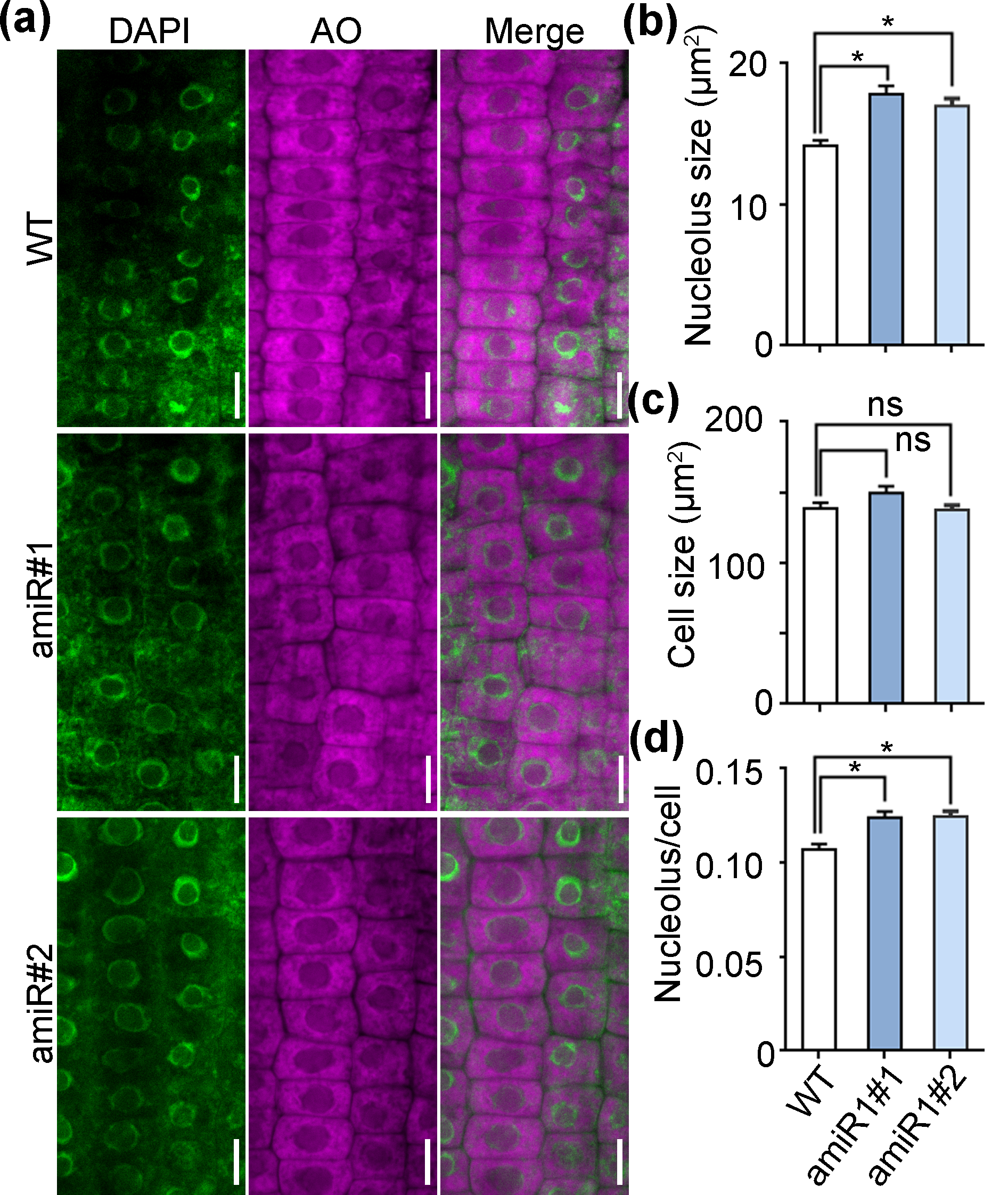

**Figure S7. *FIP1* knock-down results in enlarged nucleolus.**

(a) A representative CLSM of DAPI and acridine orange (AO) staining from wild-type or two independent transgenic lines of *amiR-FIP1* roots at 5 DAG. DAPI (green) was used to label cell nucleus, AO (magenta) was used to label nucleolus. (b-d) Quantification of nucleolus size (b), cell size (c) or ratio of nucleolus size versus cell size (d) from wild-type or two independent lines of *amiR-FIP1* roots at 5 DAG. Results are means ± standard deviation (SD, n > 30). Asterisks indicate significant difference (*t*-test, P < 0.01); ns indicates no significant differences (*t*-test, P > 0.05). Bars = 10 µm.

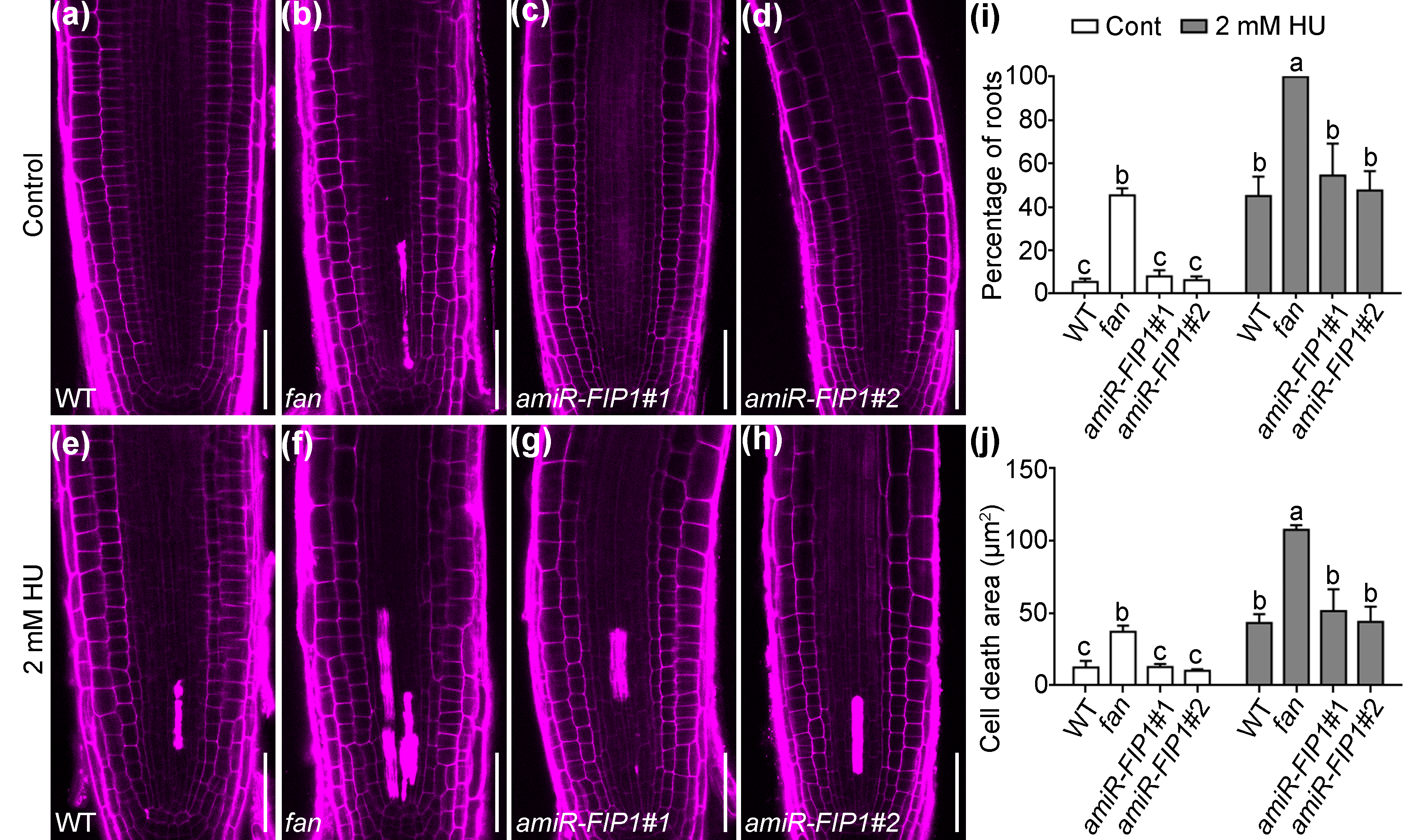

**Figure S8. *FIP1* downregulation does not show enhanced DNA damage.**

(a-h) CLSM of a PI-stained root tip from 5 DAG wild-type (a, e), *fan* (b, f), or two independent lines of *amiR-FIP1* transgenic plants (c-d, g-h) without treatment (a-d) or treated with 2 mM HU (e-h). Bars = 50 µm. (i-j) Percentage of root areas showing cell death (i) or quantification of cell death area (j). Results shown are means ± SE (n ≥ 3). In (i-j), different letters indicate significantly different groups (OneWay ANOVA, Tukey’s multiple comparisons test, P < 0.01).
