## Supplemental Table 1 for "The nucleolar complex FAN-FIP1 mediates ribosome biogenesis in Arabidopsis and is critical for BR signaling and heat tolerance"

**Table S1. Oligos used in this study.**

| Application | | No. | 5’-3’ sequences |
| --- | --- | --- | --- |
| Sequencing | *fan* | WYN-76 | GCATGATTGACTTCATTTGGAGACC |
|  |  | WYN-77 | GCTTCCTGAGGGACCTGTGGAT |
| Cloning | *FAN* CDS | P306 | TTTAAGAAGGAGCCCTTCACCATGGCTGGGGGGTCAAAAGAG |
|  |  | P307 | GGGTCGGCGCGCCCACCCTTTTAAGCTTCAGACTGAACGTTTCTG |
|  | *FAN*  (1-813bp) | P306 | TTTAAGAAGGAGCCCTTCACCATGGCTGGGGGGTCAAAAGAG |
|  |  | P1956 | GGGTCGGCGCGCCCACCCTTTCATTTTCTCTGCCATTTGTCCAC |
|  | *AATF*  (424-813bp) | P1955 | TTTAAGAAGGAGCCCTTCACCATGGATGCTGTTAAAttGGTCAAGCG |
|  |  | P1956 | GGGTCGGCGCGCCCACCCTTTCATTTTCTCTGCCATTTGTCCAC |
|  | *TRAUB*  (1048-1275bp) | P904 | TTTAAGAAGGAGCCCTTCACCATGCAAGAAGAAGGAGACCCTGAAC |
|  |  | WYN-84 | GGGTCGGCGCGCCCACCCTTTCAGAAGAGATTCTTAAGCAAATCCGCCG |
|  | *FIP1* CDS | P196 | CACCATGGAGTCACCGCACTCGAA |
|  |  | P197 | TCAGTGCCTTGTCTTTCTTTTC |
|  | *Sas10*  (70-312bp) | P1958 | TTTAAGAAGGAGCCCTTCACCATGTTGAGAGAAATGAAGAATGTTCTAG |
|  |  | P1959 | GGGTCGGCGCGCCCACCCTTTCACTTTTCCAGAAACATACGTATCTC |
|  | BRI1 CDS | ZP765 | TTTAAGAAGGAGCCCTTCACCATGAAGACTTTTTCAAGCTTCTTTCTC |
|  |  | ZP766 | GGGTCGGCGCGCCCACCCTTTAATTTTCCTTCAGGAACTTCTTTTATACTC |
|  | uORF-BRI1 CDS | WYN-354 | TTTAAGAAGGAGCCCTTCACCCTTGCATTAATTAACTTTTTTTTTATAACC |
|  |  | WYN-408 | GGGTCGGCGCGCCCACCCTTTAATTTTCCTTCAGGAACTTCTTTTATACT |
|  | *proFIP1* | P386 | TTTAAGAAGGAGCCCTTCACCAGAGAGAGGCTTTCGATTAATTTTAC |
|  |  | P387 | GGGTCGGCGCGCCCACCCTTGATTCTTGTTCAGTACTTGACAGAAG |
|  | *pro35S:ARF3-RFP* | P1935 | AGGCATGCGACGTCGGGCCCGACTAGAGCCAAGCTGATCTCC |
|  |  | P1936 | GGGGATCCTCTAGAGGGCCCTTAGGCGCCGGTGGAGTG |
| *UBQ10:amiR-FIP1* | *amiR-FIP1-pRS300* | ZP6117 | CTGCAAGGCGATTAAGTTGGGTAAC |
|  |  | ZP6118 | GCGGATAACAATTTCACACAGGAAACAG |
|  |  | P1750 | GATAAGTAACAGATGCTTGGCCCTCTCTCTTTTTGTATTCC |
|  |  | P1751 | GGAGGGCCAAGCATCTGTTACTTATTCAAAGAGAATTCAATTGA |
|  |  | P1752 | GAGGACCAAGCATCTCTTACTTTTCACGGAATCGTGATATG |
|  |  | P1753 | GAAAAGTAAGAGATGCTTGGTCCTTCTACAATATATAATTCCT |
|  | *amiR-FIP1-TOPO* | P350 | TTTAAGAAGGAGCCCTTTCACCCCAAACACACGCTCGGACGCATA |
|  |  | P351 | GGGTCGGCGCGCCCACCCTTACGAAGGCAGCATATATGTCACT |
| RT-qPCR | *FIP1* | P395 | GAAAGATGTGTTGGACGATCTG |
|  |  | P396 | AGCGCTTCTCTCTCTTCTTATC |
|  | *45S-preRNA* | P937 | GGGGGGTGGGTGTTGAGGGAG |
|  |  | P938 | GAAAAAGGGGGTTCCCACGGAC |
|  | *5’ETS1* | P1090 | CCTTGCTCGCATTGGTGAAT |
|  |  | P1091 | CGTCGACAACTTTTCCGCAT |
|  | *ITS1* | P941 | CGCGAACCAAAGATCACCA |
|  |  | P942 | GGCAAGGAATCGGCTAAGAAA |
|  | *18S rRNA* | P945 | TGACGGAGAATTAGGGTTC |
|  |  | P946 | CCTCCAATGGATCCTCGTTA |
|  | *5.8S rRNA* | P1088 | GCAACGGATATCTCGGCTCTC |
|  |  | P1089 | TGCGTTCAAAGACTCGATGG |
|  | *25S rRNA* | P5165 | CTACCGTGCGCTGGATTATGA |
|  |  | P5166 | GCTTCTAGCCCGGATTCTGACT |
|  | *EF-1α* | P982 | CTAAGGATGGTCAGACCCG |
|  |  | P983 | CTTCAGGTATGAAGACACC |
|  | *GAPDH* | ZP687 | TGAAATCAAAAAGCTATCAAGG |
|  |  | ZP688 | CATCATCCTCGGTGTATCCAA |
|  | *BRI1* | WYN-406 | CACGGCGGAAGATTGCGATA |
|  |  | WYN-407 | ATCGCACTCATCAGCCTCGC |
|  | *SAUR‐AC1* | NKP240 | AAGAGGATTCATGGCGGTCTATG |
|  |  | NKP241 | GTATTGTTAAGCCGCCCATTGG |
|  | *BR6ox1* | WYN-427 | ACGAACCACTCGGTCTTGAGGA |
|  |  | WYN-428 | TGTGACTCCAAGCTCTTCTTCATCC |
|  | *ROT3* | WYN-429 | CTCGTGGAGGAGAATATGGAGATGA |
|  |  | WYN-430 | GTCTTCATCCATGTGAACCGATATG |
| Northern blot | *18S rRNA* | WYN-618 | CATATGACTACTGGCAGGATCAACC |
|  | *5.8S rRNA* | WYN-619 | GATTCTGCAATTCACACCAAGTATC |
|  | *25S rRNA* | WYN-620 | CTCCGCTTATTGATATGCTTAAAC |
|  | *U2 snRNA* | WYN-621 | AATAGAGTTAATATCGTGTGGG |
